# Rab27-dependent egress of SARS-CoV-2 via secretory amphisomes

**DOI:** 10.64898/2025.12.15.694351

**Authors:** Wilco Nijenhuis, Hugo G.J. Damstra, Emma J. van Grinsven, Patrique Praest, Mariëlle M.P. van Grinsven, Zahra E. Soltani, Dorien C.M. de Jong, Jori Symons, Gimano D. Amatngalim, Frederik J. Verweij, Anna Akhmanova, Monique Nijhuis, Jeffrey M. Beekman, Robert Jan Lebbink, Lukas C. Kapitein

## Abstract

Severe acute respiratory syndrome coronavirus 2 (SARS-CoV-2), the causative agent of COVID-19, hijacks host cellular machinery to replicate and spread. Understanding how SARS-CoV-2 reorganizes host cell architecture to accommodate this is essential for elucidating its pathogenesis and identifying therapeutic targets. While the molecular mechanisms of SARS-CoV-2 entry are well characterized, the pathways governing viral egress remain incompletely understood. Conventional approaches such as transcriptomics and electron microscopy have provided valuable insights but lack the combined spatial and molecular resolution needed to map these processes within intact cells. Here, we apply Ten-fold Robust Expansion Microscopy (TREx) to visualize SARS-CoV-2–induced remodeling of the endolysosomal system in multiciliated cells of primary human airway epithelial tissue. This approach reveals Golgi fragmentation and the formation of enlarged virus-containing organelles. Analysis of endolysosomal markers in Vero E6 cells shows that these structures are positive for CD63, Rab7, and LC3, consistent with amphisome identity. Moreover, pharmacological inhibition of Rab27-dependent amphisome–plasma membrane fusion with Nexinhib20 reduces viral infection, implicating secretory autophagy as a pathway for SARS-CoV-2 egress. These findings establish expansion microscopy as a powerful tool for spatial virology and uncover a Rab27-mediated amphisome fusion mechanism as a druggable route for SARS-CoV-2 release.

## Introduction

COVID-19 is a respiratory illness caused by infection with severe acute respiratory syndrome coronavirus2 (SARS-CoV-2) (Steiner, Kratzel et al. 2024). After its emergence in late 2019, the SARS-CoV-2 outbreak caused a global pandemic with severe socioeconomic and medical impact, causing over 6.9 million deaths (WHO 2023). While the rollout of an effective vaccination campaign has relieved pressure on global health systems, better understanding of the SARS-CoV-2 life cycle is needed to identify novel targets to expand pharmacotherapeutic options for the treatment of COVID-19.

SARS-CoV-2 entry, replication and assembly are characterized in great detail (Steiner, Kratzel et al. 2024). During infection, viral entry is mediated by the binding of spike to the human host receptor angiotensin-converting enzyme II (ACE2) and subsequent proteolytic activation of spike by host proteases, including type II transmembrane serine protease (TMPRSS2). After entry, extensive ultrastructural rearrangements involving multiple organelles including the endoplasmic reticulum (ER) and the Golgi apparatus lead to the formation of the viral replication organelle where viral RNAs are replicated. Meanwhile, viral membrane proteins are synthesized on the ER, incorporated into transport vesicles at ER exit sites and trafficked to the endoplasmic-reticulum-Golgi intermediate compartment (ERGIC) where viral particles bud into the lumen of the ER/ERGIC and traffic to the Golgi apparatus (Pinto, Rai et al. 2022).

How viral particles traffic from the Golgi into secretory compartments is still incompletely understood (Marano, Vlachova et al. 2024, Sergio, Ricciardi et al. 2024). Originally thought to use the biosynthetic pathway for egress, more recent work reported that β-coronaviruses, including SARS-CoV-2, employ exocytic lysosomes for exit the cell (Ghosh, Dellibovi-Ragheb et al. 2020). Specifically, viral particles have been observed to localize to LAMP-1 labeled lysosomes, viral membrane proteins colocalize with lysosomal markers and β-coronavirus egress has been shown to be Rab7- and Arl8b-dependent (Ghosh, Dellibovi-Ragheb et al. 2020, Scherer, Mascheroni et al. 2022). Strikingly, lysosomes are deacidified in SARS-CoV-2 infected cells, which likely protects viral particles from lysosomal hydrolase activity (Ghosh, Dellibovi-Ragheb et al. 2020). This deacidification, and the accompanied loss of lysosomal degradative capacity, is caused by the viral ORF3A protein, which recruits and sequesters the HOPS complex, thereby blocking the fusion of Rab7-positive endosomes and autophagosomes with lysosomes (Ghosh, Dellibovi-Ragheb et al. 2020, Miao, Zhao et al. 2021, Shariq, Malik et al. 2023). It is still unknown how newly generated viral particles are transported into exocytic lysosomes, but the ORF3A-dependent block is likely to prevent the potential delivery of virions from late endosomes/multivesicular bodies to lysosomes.

Additionally, SARS-CoV-2 virions have been detected in amphisomal/endosomal structures at the early stages of infection, which has been interpreted as virions exploiting endocytosis as an additional entry route (Miao, Zhao et al. 2021). Here ORF3A is likely also involved, as both ORF3A overexpression and SARS-CoV-2 infection blocks autophagy, which results in the accumulation of autophagosomes/amphisomes (Miao, Zhao et al. 2021). However, delineation of the role of the endolysosomal system in infected cells is challenging because the system is in constant flux and cellular trafficking pathways are strongly perturbed (Sergio, Ricciardi et al. 2024). This makes it difficult to classify structures based on morphology alone, which is compounded by the fact that viral infection of individual cells is highly variable (Akilesh, Nicosia et al. 2021, Bullock, Goldsmith et al. 2021, Bullock, Goldsmith et al. 2021).

While electron microscopy (EM) is the gold standard to visualize the intracellular replication cycle in individual cells (Wolff and Bárcena, 2021), and despite advances such as correlative light and electron microscopy and 3D electron tomography (Boer et al., 2015; Lidke and Lidke, 2012), light microscopy is particularly suited to bridge scales between bulk measurements of viral replication and visualization of ultrastructure of individual cells with EM. In recent years, expansion microscopy (ExM) has emerged as a technique that enables three-dimensional super resolution imaging of biological samples with conventional microscopes. In ExM, specimens are embedded in a swellable polymer gel to physically expand the sample and isotropically increase resolution (Chen et al., 2015). As ExM relies on physical separation of fluorophores to increase resolution, this property can be exploited to visualize cellular ultrastucture using aspecific labelling to reveal local protein densities (M’Saad and Bewersdorf, 2020).

Here, we used the ExM variant TREx (Ten-fold Robust Expansion Microscopy) to visualize 3D ultrastructure in combination with labeling of viral markers in human airway cells infected with SARS-CoV-2. Upon infection with SARS-CoV-2, we observed appearance of enlarged organelles with heterogenous content that stained positive for the viral spike protein and CD63, a marker for multivesicular bodies (MVBs). Further characterization of these organelles in Vero cells indicated that these organelles are secretory amphisomes that have both endolysosomal and autophagosomal properties and depend on Rab27A for secretion. Importantly, inhibition of Rab27A-dependent secretion using Nexinhib20 reduces SARS-CoV-2 release and spreading. Together, this work underscores the power of ExM to visualize SARS-CoV-2-induced cytopathic effects in primary cell culture models and identifies a druggable pathway for viral egress.

## Results

### Expansion microscopy reveals enlarged CD63-positive late endosomal structures in SARS-CoV-2 infected human airway cells

As the upper respiratory system is the primary site of droplet-based infection by airborne viruses, including SARS-CoV-2, we adopted a recently developed protocol to differentiate human nasal epithelial cells (HNECs) from nose brushes of healthy donors (Fig. 1A) (Amatngalim, Rodenburg et al. 2022). As ciliated cells are the main permissive cell type for SARS-CoV-2 in the respiratory epithelium (Ahn, Kim et al. 2021, Pinto, Rai et al. 2022), we chose a differentiation protocol that enriched for ciliated cells. We validated the composition of our cultures by staining with cell type specific markers (Fig. S1A-B) and found a mix of basal cells (∼20-30% of total cell population), MUC5AC^+^ goblet cells (∼10-20%), ciliated cells (∼40-50%) and sparse CC10^+^ club-like cells (Fig. S1C).

**FIGURE 1:**
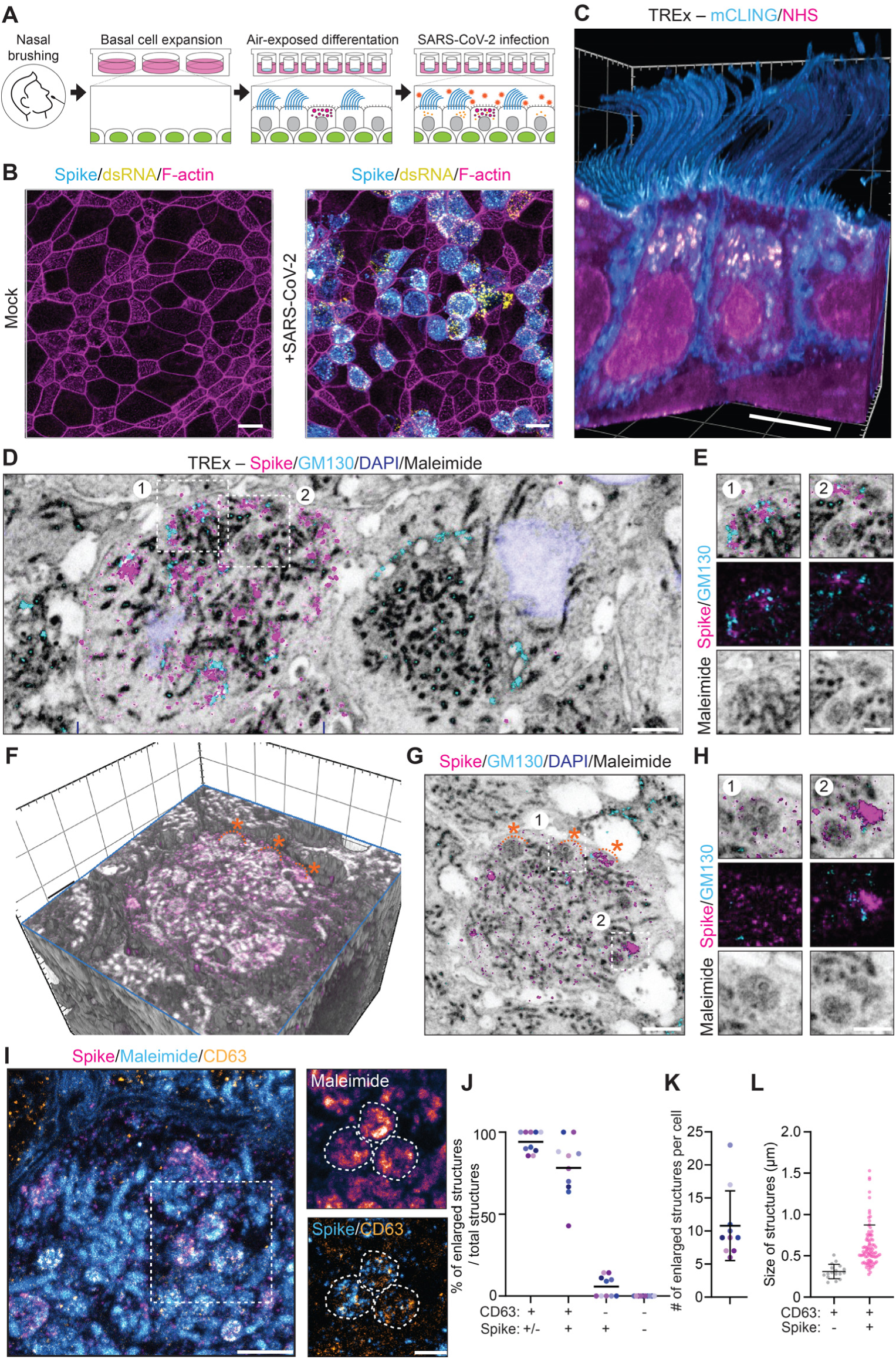
Optical nanoscopy of SARS-CoV-2 infected HNECs reveals enlarged late endosomal structures. (A) Human nasal epithelial cell (HNEC) culture system and SARS-CoV-2 infection model. Basal cells (green) are seeded and expanded. Then cells are differentiated into secretory cells (magenta) and multiciliated cell (blue) on air-liquid interface (ALI), and inoculated apically with SARS-CoV-2 virus (orange). (B) Immunofluorescent confocal images of uninfected (Mock) and SARS-CoV-2 infected (+ SARS-CoV-2) HNECs stained for spike (blue), double stranded (ds) RNA (yellow) and F-actin (Phalloidin, magenta). (C) Volumetric 3D render of TREx image of multiciliated cells stained for mCLING (total membrane, blue) and a NHS-ester (total protein, magenta). The 3D render is clipped to reveal intracellular ultrastructure. See also movie S1. (D-H) TREx images of SARS-CoV-2 infected multiciliated cells stained for spike (magenta), GM130 (blue), nuclei (DAPI, dark blue) and maleimide (total protein, black). (D and E) Maximum projection revealing ultrastructural reorganizations in a SARS-CoV-2 infected multiciliated cell. (E) Representative zooms show dispersed Golgi apparatus (E-1) and large, round, densely-stained virus-induced organelles with irregular morphology (E-2). (F) Volumetric 3D render of TREx image of a SARS-CoV-2 infected multiciliated cell and single plane corresponding to the cross-section shown (G). Orange asterisks indicate morphologically similar reorganizations. (H) Representative zooms show Spike-negative (H-1) and Spike-positive structures (H-2), resembling organelles shown in zoom E-2. See also movie S2. (I) TREx image of SARS-CoV-2 infected multiciliated cells stained for spike (magenta), CD63 (intralumenal vesicles, yellow) and maleimide (total protein, blue). Zoom shows accumulation of Spike in late endosomal compartments containing CD63-positive intralumenal vesicles. Dotted lines indicate vesicles. (J-L) Quantification of large virus-induced organelles (percentage of enlarged structures positive for CD63 and/or spike (J), number of enlarged structures per cell (K) and size of structures (L). n=number of analyzed organelles: n=108, in 10 cells, 2 independent replicates. All SARS-CoV-2 infections were performed at Multiplicity of Infection (MOI): 0.1, and cells were fixed at 3 days post infection (dpi). Scale bars are 10 µm (B), ∼2 μm (C), ∼1 μm (D,F,G,I), or 500 nm (all zooms). TREx scale bars were corrected to indicate pre-expansion dimensions.

Next, we set out to validate our infection model. Fully differentiated HNEC cultures were inoculated apically with SARS-CoV-2 from a clinical isolate (NL/2020) at a Multiplicity of Infection (MOI) of 0.1 for 2h, before washing and incubating for up to 3 days post infection (dpi) before fixation. At 3 dpi, patches of infected cells could be found among regions devoid of infected cells (Fig. 1B). By correlating spike expression with cell type-specific markers, we found that ciliated cells were preferentially infected in our infection model (Fig. S2D), which is in line with other reports (Ahn, Kim et al. 2021, Pinto, Rai et al. 2022).

To visualize cellular remodeling in SARS-CoV-2 infected cells at high resolution, we set out to visualize HNEC ultrastructure after infection combined with molecular specificity. Using an optimized TREx protocol (Damstra, Mohar et al. 2022, van Grinsven, Katrukha et al. 2025), we could expand HNEC cultures directly from Transwell filters and readily visualize cellular ultrastructure. We labeled cellular membranes using mCLING, a palmitoylated small peptide that intercalates into the lipid bilayer and is compatible with TREx chemistry (Revelo, Kamin et al. 2014, Damstra, Mohar et al. 2022), and combined it with a fluorophore-labeled N-hydroxysuccinimide (NHS) ester or maleimide that covalently binds to lysines or cysteines, respectively (Fig. 1C, S2A) (M’Saad and Bewersdorf 2020). Optical sectioning revealed a variety of structures with a contrast reminiscent of electron microscopy, including extensive interdigitated membrane contacts between adjacent cells (Fig. S2C), and protein-dense complexes such as basal bodies at the base of cilia that could be morphologically identified (Fig. S2D, S2E).

To further explore the intracellular organization of infected HNECs, we used a general protein stain in combination with the selective labeling of the viral protein spike and GM130, a marker for cis-Golgi (Fig. 1D). Uninfected cells revealed perinuclear ribbon-like Golgi structures (Fig. S2F). In infected cells, we observed fragmentation of the Golgi ribbons, consistent with previous reports (Scherer, Mascheroni et al. 2022, Zhang, Kennedy et al. 2024), with the Golgi fragments often neighbored by spike-positive densities (Fig. 1E, S2F). In addition, we observed actin remodeling in SARS-CoV-2 infected cells, including the formation of numerous spike-positive branched microvilli and membranous structures, consistent with recent EM studies (Fig. S2G, S2H) (Pinto, Rai et al. 2022, Wu, Lidsky et al. 2023).

Strikingly, the total protein stain revealed a different type of compartment that was not observed in uninfected cells (Fig. 1F-H) These protein-dense, large, round organelles with heterogenous content were devoid of GM130 staining and typically contained various sparse spike densities. Based on the morphology and intraluminal content of these enlarged organelles, we rationalized that if these organelles were MVBs containing intraluminal vesicles (ILVs), the organelles should be decorated by the ILV marker CD63 (Fig. 1I). Indeed while 94 ± 6% (mean ± SD) of the SARS-CoV-2 induced organelles were positive for CD63 and 78 ± 18% were positive for both CD63 and spike, just 7 ± 7% were decorated only by spike (Fig. 1J). Structures that were labeled by neither CD63 nor spike were not observed (n = 108 structures from 10 cells, Fig. 1K). CD63 and spike were typically detected throughout these large compartments (Fig. 1L), but were sometimes enriched at the periphery of the largest of these structures. Moreover, the vesicles containing both CD63 and spike were on average 2 times larger than CD63-positive vesicles without spike. These results indicate that SARS-CoV-2 infection results in the formation of enlarged CD63-positive MVBs, a type of late endolysosomes.

### SARS-CoV-2 promotes the formation of secretory amphisomes

To understand the identity and function of the enlarged MVBs we observed after SARS-CoV-2 infection, we set out to systematically characterize late endolysosomal compartments in infected cells, compared to uninfected cells. To reduce complexity, we first moved our infection system to Vero E6 cells, a highly permissive cell type that lacks an interferon response and is commonly used to study viral infection, including SARS-CoV-2 (Ng, Tan et al. 2003). Quantitative analysis of viral markers including spike and double stranded (ds) RNA, a marker for viral replication organelles, revealed a progressive increase of viral markers with increasing severity of infection at a cellular level (S3A-C), in line with previous reports (Scherer, Mascheroni et al. 2022), and recapitulated key observations from expansion microscopy, including Golgi fragmentation (Fig. S2F, S3A, S3D). Moreover, we confirmed the presence of enlarged CD63 and spike-positive endolysosomes in infected Vero E6 cells with TREx (Fig. S4). While Golgi fragmentation was observed using cis-Golgi marker GM130, we found a near complete loss of the trans-Golgi marker TGN46 even in early stages of infection, i.e. low spike levels and non-fragmented cis-Golgi, suggesting a gross perturbation of the trans-Golgi in SARS-CoV-2 infected cells (Fig. S3E-H). Moreover, this is consistent with the notion that viral egress is independent of the biosynthetic pathway (Ghosh, Dellibovi-Ragheb et al. 2020) .

Next, to probe the late endolysosomal system in infected cells, we stained cells for the late endosomal/MVB markers CD63 and Rab7 and the lysosomal marker LAMTOR4 (Fig. 2A-C). Late endolysosomes share many of these membrane markers and undergo continuous maturation. Under basal conditions, the late endolysosomal markers CD63, Rab7, and LAMTOR4 show a punctate localization and this puncta are distributed throughout the cell. In SARS-CoV-2 infected cells, we observed the formation of enlarged endolysosomal vesicles, which were enriched in the perinuclear region and decorated by CD63 and Rab7. Strikingly, whereas LAMTOR4 was present on late endolysosomal vesicles, it was strongly depleted from enlarged organelles in SARS-CoV-2 infected cells (Fig. 2C), suggesting that lysosomal function and identity is lost after infection. Since SARS-CoV-2 has been demonstrated to block HOPS-dependent fusion of MVBs with lysosomes via ORF3A (Miao, Zhao et al. 2021, Zhang, Sun et al. 2021) we reasoned that the depletion of LAMTOR4 from organelles could be caused in part by impeded endolysosomal maturation. Indeed, whereas CD63 and LAMTOR4 mostly stained the same organelles under basal conditions, these markers decorated more distinct populations in infected cells (Fig. 2D, 2E), suggesting a block in endolysosomal maturation from late endosomes to lysosomes. Consistently, lysosome deacidification, a hallmark of lysosome dysfunction, has been reported after SARS-CoV-2 infection (Ghosh, Dellibovi-Ragheb et al. 2020), and labeling of markers of the mannose-6-phosphate pathway, M6PR and Clathrin Heavy chain revealed a near complete loss of M6PR from the Golgi apparatus, suggesting disrupted trafficking through the mannose-6-phosphate pathway (Fig. S3I).

**FIGURE 2:**
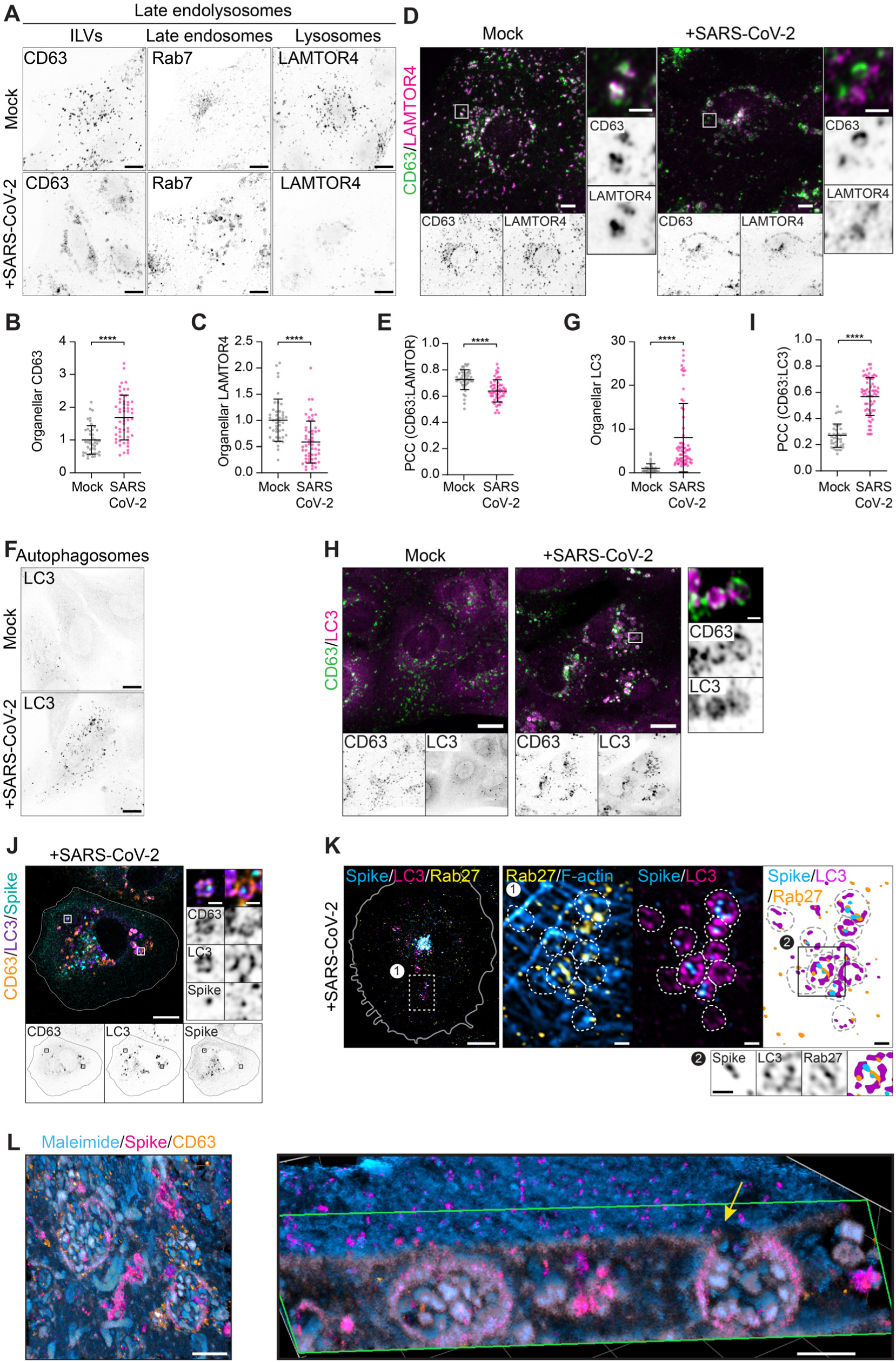
SARS-CoV-2 induces the formation of Rab27-positive secretory amphisomes. (A-K) Representative confocal images (A, D, F, H, J, K) and quantification (B, C, E, G, I) of uninfected (Mock) and SARS-CoV-2 infected (+ SARS-CoV-2) Vero E6 cells stained for the following organellar markers; (A-left) CD63 (intraluminal vesicles (ILVs), black); (A-middle) Rab7 (late endosomes, black); (A-right) LAMTOR4 (lysosomes, black); (D) CD63 (green) and LAMTOR4 (magenta) where zooms show (absence of) colocalization of CD63 and LAMTOR4; (F) LC3 (autophagasomes, black); (H) CD63 (green) and LC3 (magenta) where zoom shows colocalization of CD63 and LC3; (J) CD63 (yellow), LC3 (purple) and spike (blue) where zooms show amphisomes with Spike-positive puncta; (K) spike (blue), LC3 (magenta) and Rab27 (yellow) where zooms show enlarged spike- and Rab27-positive vesicles (K-1) and single vesicle (K-2). The corresponding quantifications in (B), (C) and (G) show the normalized organelle marker intensity (defined as (mean organellar intensity – mean non organellar intensity) x organelle area) per cell. The graphs in (E) and (I) show the non-thresholded Pearsons correlation coefficient (PCC). Dots represent one cell, bars indicate mean±SD. n=number of analyzed cells: n=37-64 per condition, 3 independent replicates (refer to supplemental table 1 for details). Asterisks indicate significance (Student’s *t* test, unpaired). ∗∗∗∗, P *<* 0.0001. (L) TREx imaging (left) of SARS-CoV-2 infected Vero E6 cells stained for spike (magenta), CD63 (yellow) and maleimide (total protein, blue) showing enlarged Spike-positive amphisome. Volumetrically rendered side-view (right), clipped to reveal one enlarged virus-containing amphisome continuous with the plasma membrane (arrow). See also movie S3. Scale bars are 10 µm (A, D, F, H, J, K) 1 µm (Zooms), ∼2 µm (L). TREx scale bars were corrected to indicate pre-expansion dimensions. All SARS-CoV-2 infections were performed at Multiplicity of Infection (MOI): 0.01, and cells were fixed at 2 days post infection (dpi).

When we examined autophagosomes by staining for LC3, we observed a strong induction of the autophagy pathway in SARS-CoV-2 infected cells, including the formation of large perinuclear LC3-positive vesicles (Fig. 2F, 2G). Since SARS-CoV-2 has been detected in both the autophagosomal and endolysosomal systems, and since autophagosomal secretion is dependent on fusion with endolysosomes to obtain the required membrane machinery for tethering and fusion with the plasma membrane (Ponpuak, Mandell et al. 2015, Kimura, Jia et al. 2017, Ganesan and Cai 2021), we reasoned that SARS-CoV-2 might utilize a hybrid of both pathways, so-called secretory amphisomes, for egress. Secretory amphisomes are non-degradative organelles that form an unconventional secretion pathway that arises through the fusion of endosomes with autophagosomes, and are decorated by CD63 and LC3 (Ganesan and Cai 2021). We therefore examined the localization of CD63 and LC3 together and observed that many of the enlarged perinuclear vesicles were positive for both markers, indicating that amphisomes were indeed formed in SARS-CoV-2 infected cells (Fig. 2G). This was supported by a strong increase in colocalization between CD63 and LC3 (Fig. 2I). Moreover, sparse spike positive puncta could be observed within these amphisomes (Fig. 2J).

Next, we wondered whether the enlarged amphisomes would be decorated by the GTPase Rab27, which is a key marker of secretory endolysosomes and has also been implicated in secretory autophagy (Solvik, Nguyen et al. 2022). Indeed, when we combined labeling of spike, LC3 and Rab27, we found colocalization of all 3 markers on the same organelles (Fig. 2K), suggesting that the amphisomes that form after SARS-CoV-2 infection take on a secretory identity. Consistently, volumetric reconstruction of expanded Vero E6 cells stained for total protein, spike and CD63 revealed a putative fusion event of a spike- and CD63-positive enlarged amphisomes with the plasma membrane (Fig. 2L, Fig. S4). Thus, taken together, we find that SARS-CoV-2 infection induces the formation of enlarged secretory amphisomes, which are decorated by CD63, Rab7, Rab27 and LC3, and low levels of LAMTOR4, which contain spike-positive puncta and potentially contribute to viral egress via Rab27A-dependent secretion.

### Inhibition of Rab27A-dependent secretion blocks SARS-CoV-2 infection

Rab27 is a key player in controlling amphisome secretion by docking to the plasma membrane via its binding partner (JFC1) (Solvik, Nguyen et al. 2022). We therefore hypothesized that pharmacological inhibition of Rab27A-dependent secretion would limit SARS-CoV-2 release from infected cells, effectively trapping the virus inside. Nexinhib20 has been identified as a potent inhibitor of Rab27A-dependent secretion by preventing docking of Rab27A-positive vesicles to the plasma membrane (Fig. 3A) (Johnson, Ramadass et al. 2016). We first validated Nexinhib20 in Vero E6 cells by testing its ability to inhibit MVB secretion, a related secretory process dependent on Rab27. To this end, we performed TIRF microscopy of CD63-pHluorin, a pH-sensitive fluorophore that is quenched in the acidic lumen of MVB’s but lights up in the neutral extracellular environment upon secretion, which allowed us to directly visualize plasma membrane fusion events of MVBs (Verweij, Bebelman et al. 2018). Under basal conditions, we monitored 12 ± 4 secretion events per cell per minute, but in cells that were treated with a concentration range of Nexinhib20 secretion was blocked in a dose-dependent manner, with a potent block (3 ± 3 events per cell per minute) at 320 nM (reported IC50: 330 nM); a near complete block (0.8 ± 1.2 events per cell per minute) was achieved at 2000 nM (Fig. 3B, 3C). Whereas treatment with this highest dose of Nexinhib20 decreased cell density, indicative of impeded cell proliferation or elevated cell death, treatment with lower concentrations for up to 48 hours did not significantly affect cell viability (Fig. 3D). We reasoned that if SARS-CoV-2 egresses via secretory amphisomes in a Rab27-dependent manner, treatment with Nexinhib20 should cause the virus to accumulate in amphisomes. To test this, we treated infected Vero E6 cells with Nexinhib20 (800 nM) and visualized CD63 and spike. Indeed, treatment with Nexinhib20 caused the appearance of CD63-positive vesicles that were densely labeled with spike, with the average spike intensity in CD63-positive vesicles increasing 7-fold (7.2 ± 21.8 events per cell per minute; Fig. 3F). In control cells, we observed only sparse labeling of spike in CD63-positive vesicles, in line with continuous viral secretion, and dispersal of spike, in absence of Nexinhib20. We found a similar accumulation of spike in LC3-positive vesicles (Fig. 3G), further supporting the notion that SARS-CoV-2 accumulates in amphisomes. Finally, we tested if blocking Rab27-dependent egress would limit SARS-CoV-2 infection. Therefore, we treated infected cells with a concentration range of Nexinhib20 and directly quantified viral SARS-CoV-2 RNA levels by qRT-PCR, which again showed a dose dependent decrease in viral genome at both 24 and 48 hours post infection (Fig. 3H). Notably, treatment with 800 nM Nexinhib20 reduced viral genome levels by >900 fold at 24 hpi (9.85 ΔCT) and >400.000 fold at 48 hpi (18.67 ΔCT). While moderate concentrations of Nexinhib20 did not completely block viral spread, we did observe a delayed infection that could imply that viral release was inhibited from a large fraction of cells. Finally, when we examined viral spread within whole cultures by high content imaging (Fig. 3I), we found that treatment with both 320 nM and 800 nM Nexinhib20 reduced the fraction of infected cells at 24 hours post infection by ∼7 fold and ∼18 fold respectively (Fig. 3J). Again, Nexinhib20 treatment did not completely block SARS-CoV-2 proliferation, as the fraction of infected cells did increase at 48 hour post infection in Nexinhib20 treated cells. Together, these data indicate that SARS-CoV-2 egress is mediated by secretory amphisomes with both endolysosomal and autophagosomal properties and relies on Rab27A-dependent secretion, which can be blocked using Nexinhib20 (Fig. 4).

**FIGURE 3:**
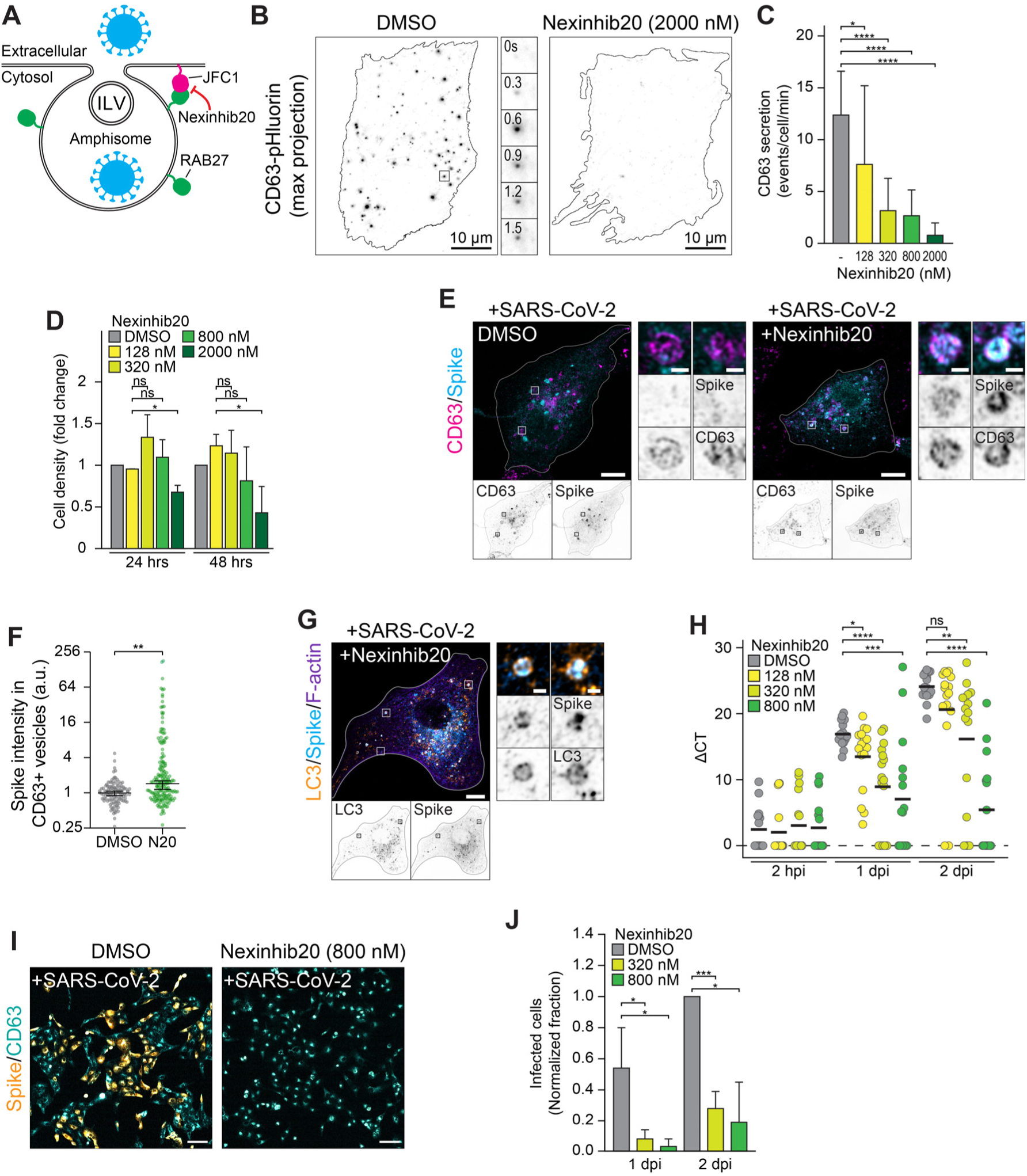
Inhibition of Rab27-dependent secretion blocks SARS-CoV-2 infection. (A) Model of Rab27-dependent SARS-CoV-2 secretion and inhibitory mechanism by Nexinhib20. Intraluminal vesicles (ILV). (B) Maximum (max) projection of total internal reflection fluorescence (TIRF) microscopy images of CD63-pHluorin secretion events in Vero E6 cells treated with DMSO (left) or 2000 nM Nexinhib20 (right). Zoom shows example of one secretion event at different timepoints. (C) Quantification of the number CD63-pHluorin secretion events per cell per minute (min) in Vero E6 cells treated with different concentrations of Nexinhib20. n=number of analyzed cells: n=14-19 per condition, 3 independent replicates (refer to supplemental table 1 for details). (D) High-content imaging-based quantification of Vero E6 cell density in the presence of different concentrations of Nexinhib20 after 24 or 48 hours compared to control, DMSO treated Vero E6 cells (e.a. a fold change of 1 indicates the cell density of the Nexinhib20 sample is similar to the DMSO treated sample). n=number of analyzed field of views: n= 4-5 FOV in independent replicates (refer to supplemental table 1 for details). (E) Vero E6 cells infected with SARS-CoV-2 and treated with DMSO (left) or 800 nM Nexinhib20 (right) and stained for CD63 (magenta) and spike (blue). Zooms show Spike-negative CD63-positive vesicles (DMSO) or Spike-positive CD63-positive vesicles (Nexinhib20). (F) Quantification of the Spike intensity in CD63-positive vesicles in SARS-CoV-2 infected Vero E6 cells normalized to the DMSO control (y axis is shown with a 4-fold increments scale). 3 independent replicates (refer to supplemental table 1 for details). (G) Vero E6 cells infected with SARS-CoV-2 and treated for with 800 nM Nexinhib20 and stained for LC3 (yellow), Spike (blue) and F-actin (phalloidin, purple). Zooms show Spike-positive LC3-positive vesicles. (H) qRT-PCR of Vero E6 cells infected with SARS-CoV-2 and treated for XX hours with indicated concentration Nexinhib20 and fixed at different timepoints (hour (h) and days (d) post infection (pi)). ΔCT is the normalized expression of the SARS-CoV-2 E gene to a reference gene. n=number of analyzed wells: n= 16 from 3 independent replicates (refer to supplemental table 1 for details). (I) High-content-based imaging of SARS-CoV-2-infected Vero E6 cells treated with DMSO or 800 nM Nexinhib20 and stained for spike (yellow) and CD63 (blue). (J) Quantification of the fraction of SARS-CoV-2 infected cells to the total number of cells on the coverslip in DMSO treated or Nexinhib20 treated cells at 1 or 2 dpi. Bar graphs represent mean±SD. In graph (H), dots represent individual wells. See also supplemental table 1. **** p<0.0001, *** p<0.001, ** p<0.01, * p<0.05, ns = non-significant (Student’s *t*-test, unpaired). Scale bars are 10 µm (B,E), 100 µm (G).

**FIGURE 4:**
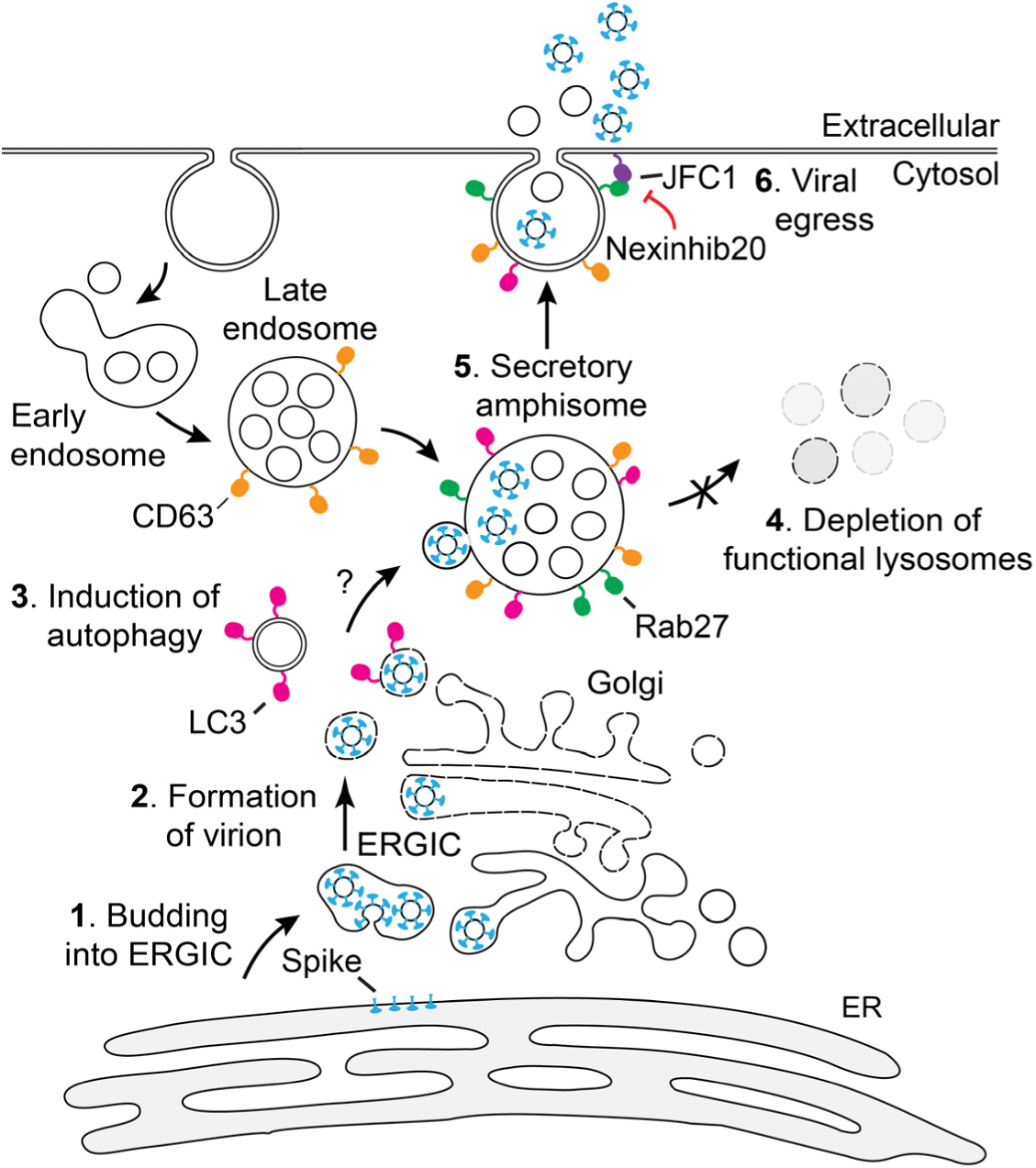
Model of SARS-CoV-2 egress via secretory amphisomes. Viral particles bud from the ERGIC (1,2) and undergo autophagy (3) or are otherwise transferred into LC3-positive carriers. ORF3A blocks the fusion of late endosomes with lysosomes (4), promoting the formation of amphisomes (5) (the result of fusion between autophagosomes and late endosomes) that can fuse with the plasma membrane in a Rab27-dependent manner (6). Nexinhib20 blocks this process and reduces the spreading of SARS-CoV-2 infection.

## Discussion

Here, we have used innovative light microscopy to visualize key aspects of the SARS-CoV-2 infectious cycle in primary human airway cells and Vero E6 cells. Since viral infection of individual cells is highly variable, we explored the use of ExM to bridge scales between bulk interrogation of viral replication and EM, which is the traditional method to visualize the intracellular viral replication cycle in individual cells (Wolff and Bárcena, 2021). Whereas multiple aspects of the viral replication cycle were already discovered using EM, ExM can complement such studies by providing both ultrastructural information and specific labeling. This in turn will help us understand how SARS-CoV-2 highjacks the endolysosomal system to its advantage. In our infection model, we observed a large-scale reorganization of cellular architecture, including Golgi fragmentation, extensive remodeling of the endolysosomal network, and the formation of enlarged secretory amphisomes. Moreover, we identified a druggable pathway for viral egress. We expect that our methodology will be a valuable addition to existing techniques for visualizing viral infection in general and help further our understanding of SARS-CoV-2 induced cytopathy.

In SARS-CoV-2 infected cells, we observed enlarged organelles that formed after infection and were decorated by viral protein spike. Based on the morphology of these organelles, we first examined whether these organelles were multivesicular bodies, but later found that these structures were positive for both CD63 and LC3 (markers for MVBs and autophagosomes, respectively). Moreover, we observed a depletion of lysosomal markers in infected cells, as well as an increase in organelle-associated LC3, consistent with previous reports (Hui et al., 2021; Li et al., 2021). Interestingly, when lysosome fusion with autophagosomes is impaired, these autophagosomes can be diverted to late endosomes and result in the formation of hybrid organelles, termed secretory amphisomes, that can fuse with the plasma membrane to release the autophagosomal content via secretion (Solvik et al., 2022). Indeed, SARS-CoV-2 blocks lysosome acidification (Ghosh et al., 2020), and the viral protein ORF3a is involved in inhibiting autophagy by preventing autophagosome fusion with lysosomes (Zhang, Sun et al. 2021, Solvik, Nguyen et al. 2022, Miller, Houlihan et al. 2023). Therefore, we propose that the enlarged organelles observed after SARS-CoV-2 infection are secretory amphisomes with both endolysosomal and autophagosomal properties.

We confirmed that these amphisomes take on a secretory identity as we found co-localization of spike-positive amphisomes, characterized by the presence of intraluminal vesicles as well as autophagy cargo receptors, with Rab27A, a GTPase involved in both ILV secretion (Ostrowski, Carmo et al. 2010) and amphisome secretion (Solvik, Nguyen et al. 2022). Consistently, β-coronaviruses, including SARS-CoV-2, do not rely on the biosynthetic secretory pathway for egress (Ghosh et al., 2020). Furthermore, enlarged late endosomal compartments after viral infection were previously observed using EM (Cortese et al., 2020; Zhou et al., 2017), SARS-CoV-2 virions have also been detected in amphisomal/endosomal structures (Miao, Zhao et al. 2021), and amphisomes have been suggested to be a route of viral egress for various viruses, including some coronaviruses (Ponpuak et al., 2015; Teo et al., 2021). Moreover, in non-infected cells, disruption of lysosome fusion with late endosomes and autophagosomes, by deletion of HOPS, BORC or Arl8b induces the formation of enlarged, non-acidified, perinuclear amphisomes and endolysosomes which are strikingly similar to those observed in SARS-CoV-2 infected cells (van der Beek, de Heus et al. 2024). This fusion block additionally induces exosome secretion and upregulated secretion of lysosomal enzymes (Shelke, Williamson et al. 2023, van der Beek, de Heus et al. 2024). The highly similar effects of SARS-CoV-2 infection, ORF3A overexpression and HOPS/BORC/Arl8b deletion supports the notion that SARS-CoV-2 reroutes the canonical degradative endolysosomal maturation pathway towards an unconventional non-degradative amphisome secretion pathway, which is utilized for egress. Importantly, we show that viral release can be delayed by treating cells with Nexinhib20, an inhibitor of Rab27A-mediated secretion in a dose-dependent response. Together, these results support a model in which the ORF3A blocks the fusion of lysosomes with late endosomes and autophagosomes, which leads to the accumulation of amphisomes that mediate the egress of SARS-CoV-2 via Rab27A-mediated secretion.

How spike is trafficked to amphisomes remains incompletely understood. As SARS-CoV-2 buds at the ERGIC, and LC3 positive precursor vesicles have been described after viral infection near the ERGIC (Reggiori, Monastyrska et al. 2010), a potential explanation of how SARS-CoV-2 gets routed to amphisomes could be via LC3-positive precursor vesicles during budding from the ERGIC. Alternatively, macroautophagy could play a role in targeting virions to amphisomes. Future experiments are needed to explore the exact interplay between these different compartments during virion formation and egress.

Our approach illustrates how innovative light microscopy, and in particular ExM, can provide novel insights into COVID-19 progression, which ultimately could help guide the development of therapeutics. For example, most work on developing antiviral agents focusses on the viral proteins such as the RNA-dependent RNA polymerase (RdRp) (e.g., Remdesivir, Favipiravir, Ribavirin, Molnupiravir), the main viral proteases (e.g., Paxlovid, Lopinavir, Ritonavir), or interference between the association of SARS-CoV-2 with the main entry receptor ACE2 or protease TMPRSS2 (e.g., Camostat mesylate). However, viral targets are highly susceptible to the high mutation rate of viruses, which limits the scope to highly conserved proteins such as the RdRp. By targeting the host cellular secretion machinery, we have identified a druggable pathway that is completely distinct from the existing SARS-CoV-2 antivirals, and is less susceptible to the mutation rate of the viral proteins albeit with a higher chance of toxicity. This enlarges the toolbox of antivirals that could be included in combinatorial treatment regimens and provides a range of druggable targets to guide drug repurposing screens and future drug development.

## Material and Methods

### Key resources table

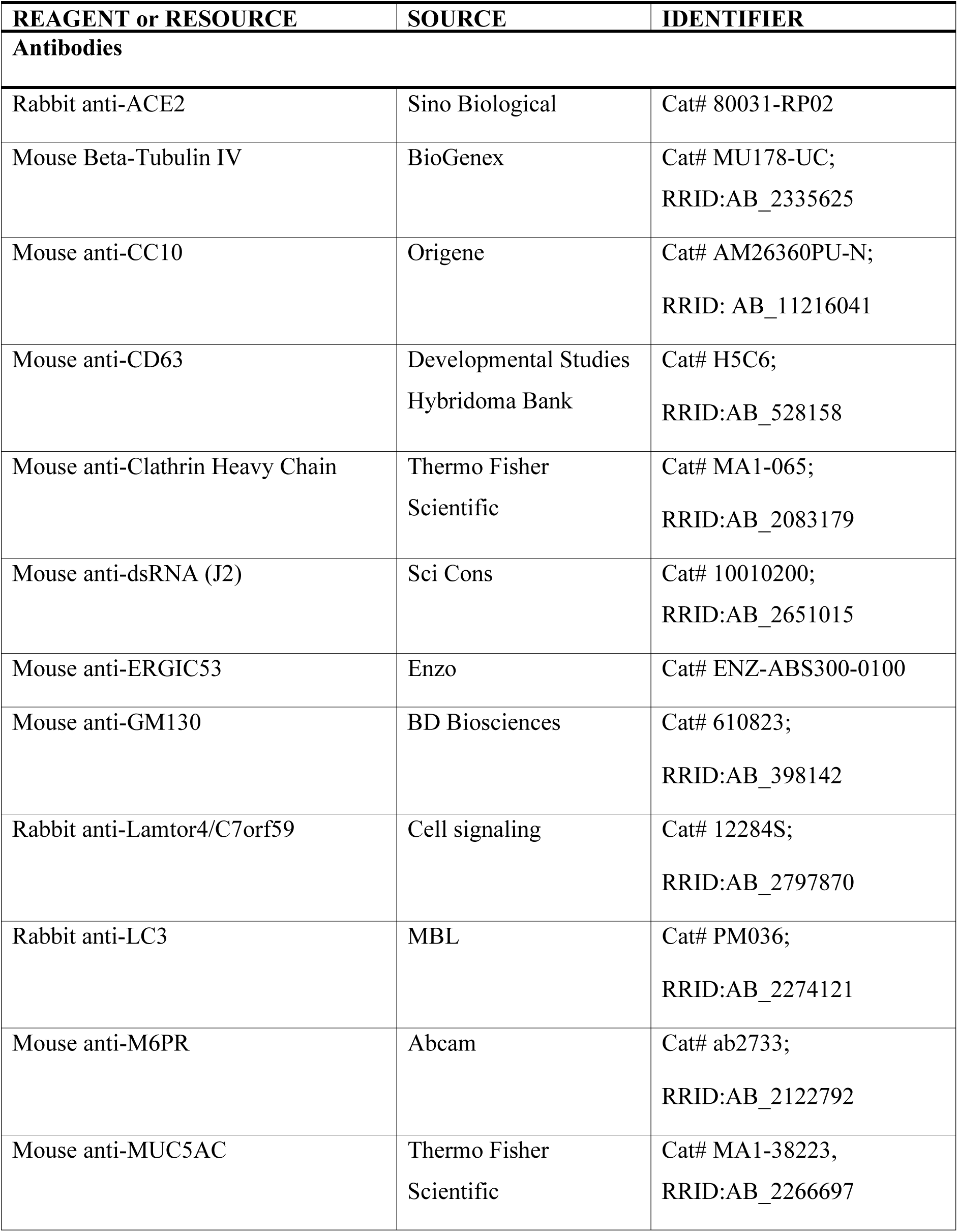
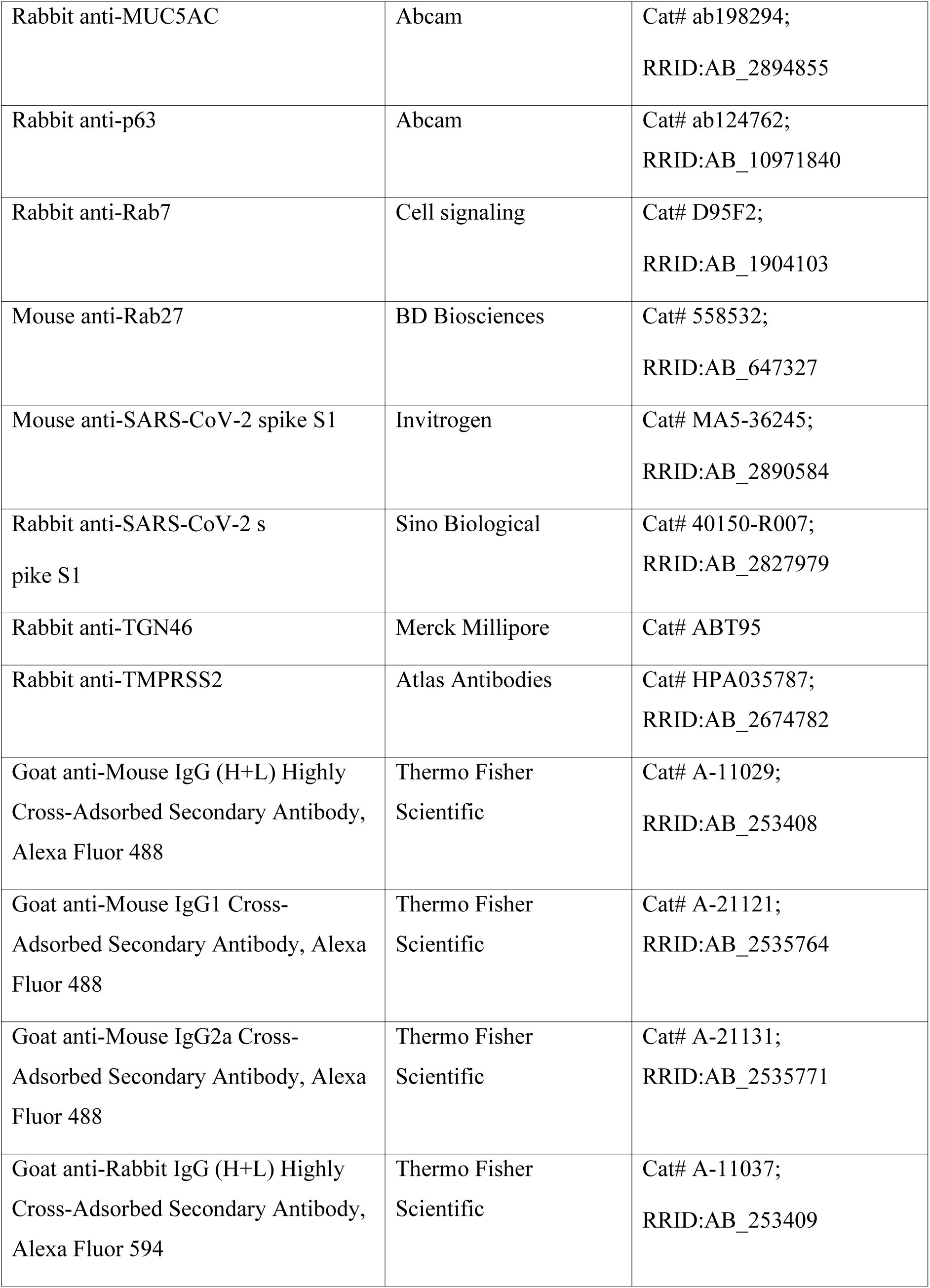
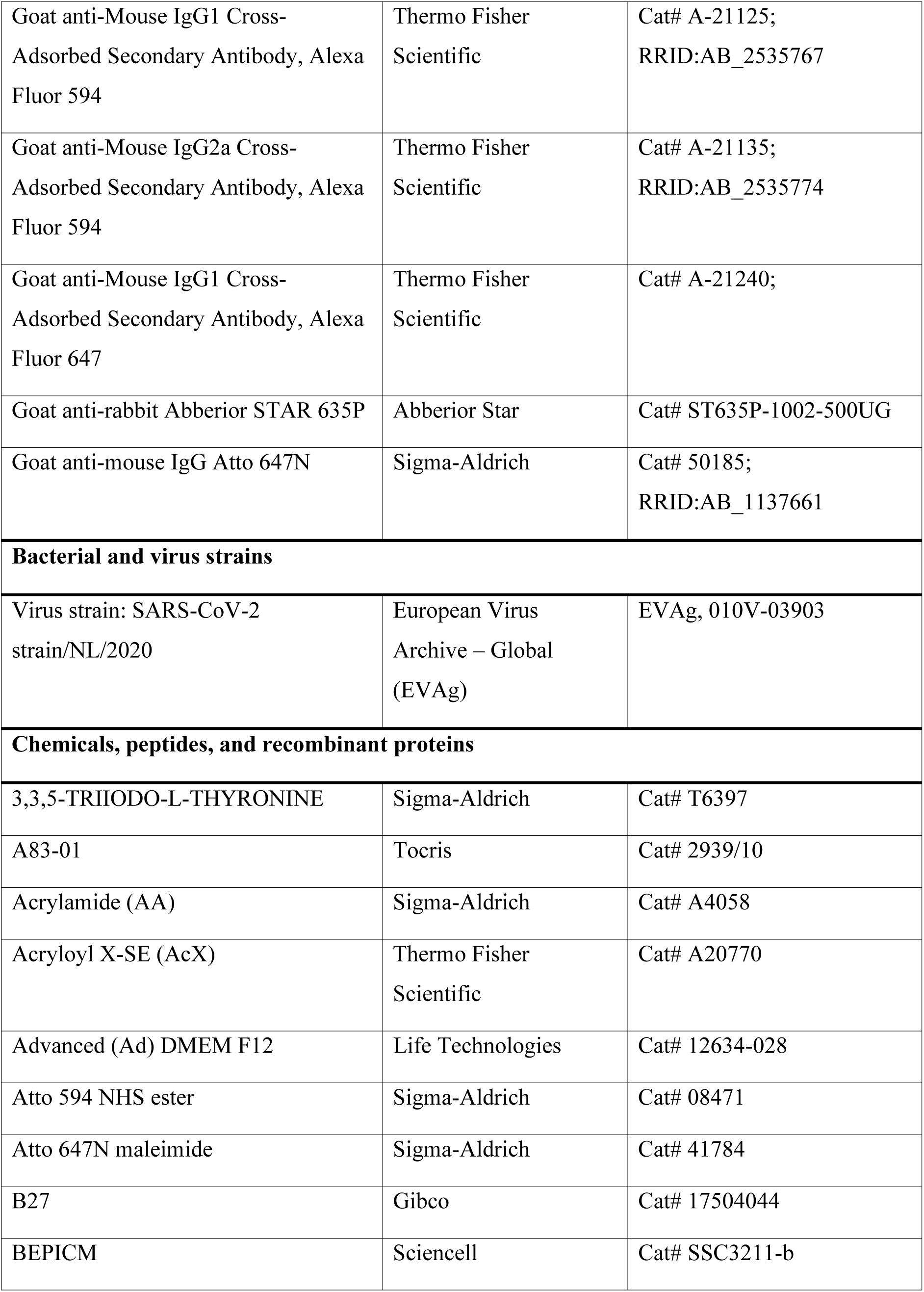
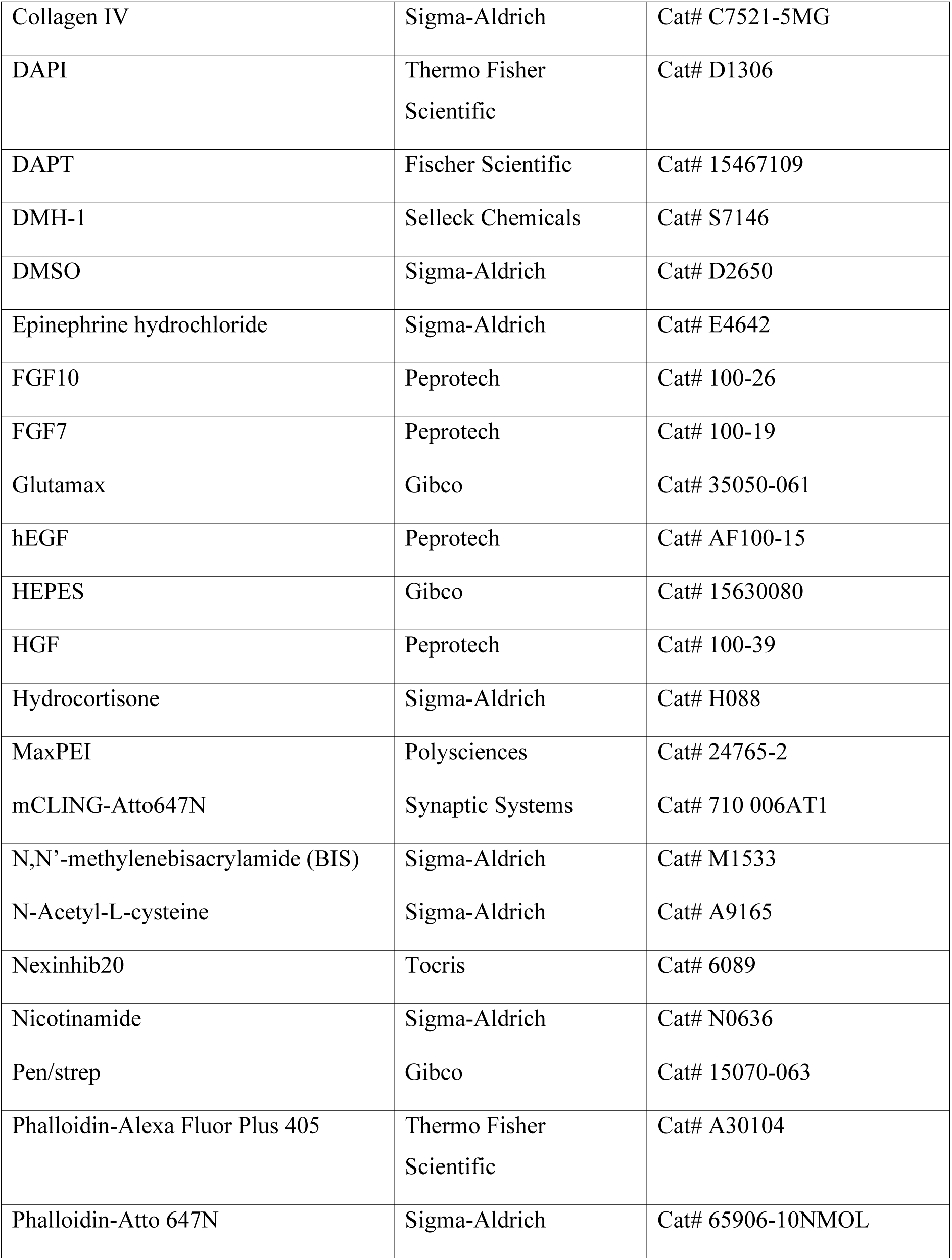
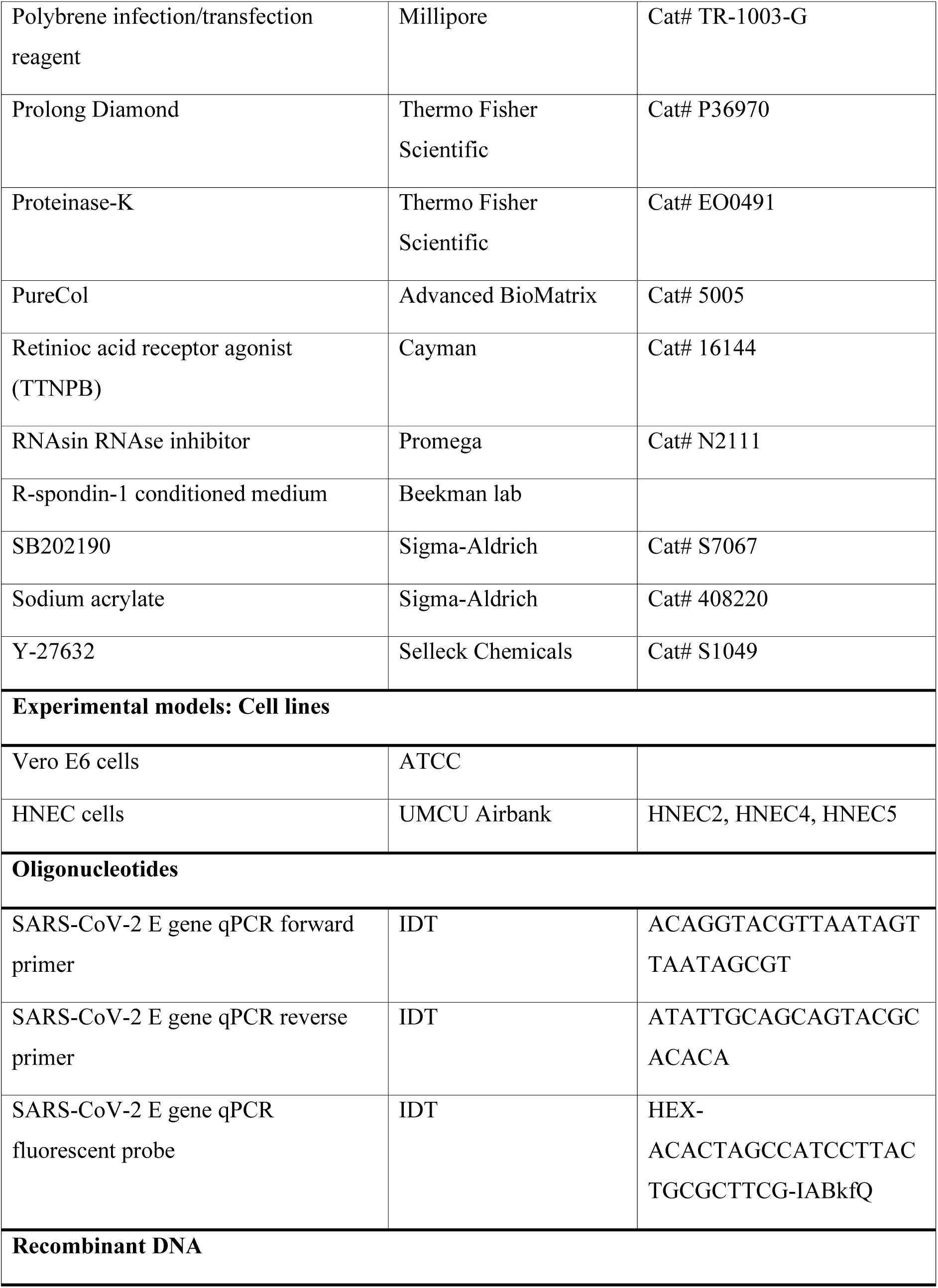
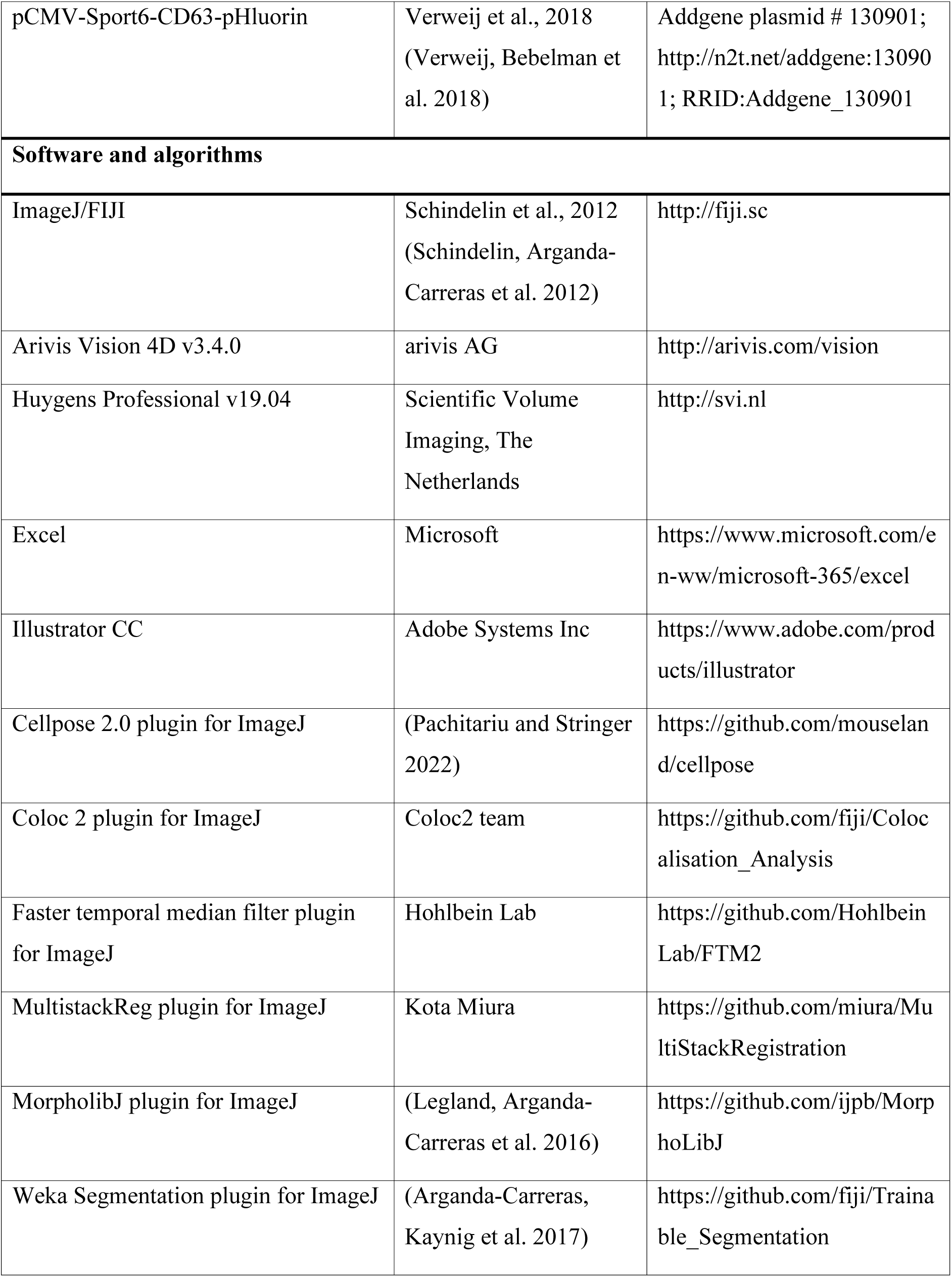
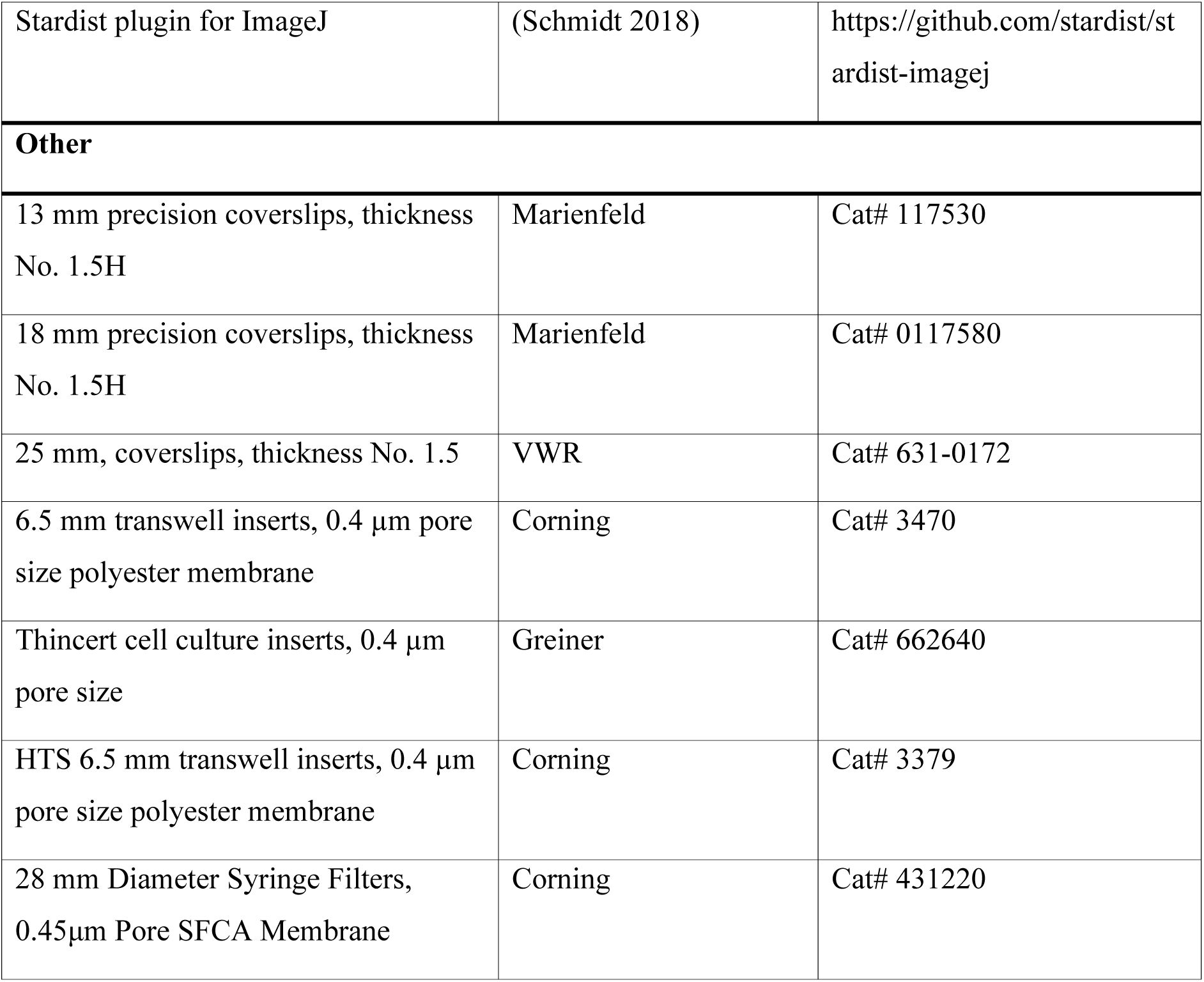

## Material and Methods

### Cell culture Vero E6 and HEK239T cells

Vero E6 cells and HEK293T cells were maintained in Dulbecco’s modified Eagle’s medium (DMEM, Invitrogen) supplemented with 10% fetal calf serum (FCS, PAA laboratories), 100 Units/ml penicillin, 100 µg/ml streptomycin and 2mM L-glutamine.

### Human airway epithelial cells culture and differentiation

Adult human airway cells were collected from nasal brushings of healthy volunteers without respiratory tract symptoms. All donors provided explicit informed consent and the study was approved by the Medical Research Ethics Committee (Tc-BIO protocols 16-856 and 20/806). The material used in this study was derived from two independent donors. All data was derived from human nasal epithelial cell (HNEC) donor HNEC2 with the exception of data shown in Figures 1C and S2, which were derived from donor HNEC5. HNECs were cultured as described previously (Amatngalim, Rodenburg et al. 2022). In brief, HNECs were expanded on collagen IV-coated (50 µg/ml, Sigma Aldrich)-6-well culture plates using basal cell (BC) expansion medium consisting of BEPICM (50%, Sciencell #3211), Advanced (Ad) DMEM F12 (23,5%, Life Technologies #12634-028), R-spondin-1 conditioned medium (20%), HEPES (10 mM, Gibco #15630080), Glutamax (1x, Gibco #35050-061), Pen/strep (1x, Gibco #15070-063), B27 (2%, Gibco #12587010), Hydrocortisone (0.5 mg/mL, Sigma #H088), 3,3,5-triiodo-L-thyronine (100 nM, Sigma #T6397), Epinephrine hydrochloride (0.5 mg/mL, Sigma #E4642), N-Acetyl-L-cysteine (1,25 mM, Sigma #A9165), Nicotinamide (5 mM, Sigma #N0636), A83-01 (1 mM, Tocris #2939/10), DMH-1 (1 µM, Sellech Chemicals, #S7146), Y-27632 (5 µM, Selleck Chemicals #S1049), SB202190 (500 nM, Sigma #S7067), HGF (25 ng/mL, Peprotech # 100-39), hEGF (5 ng/mL, Peprotech #AF100-39), DAPT (Fischer Scientific #15467109), FGF7 (25 ng/mL, Peprotech #100-19) and FGF10 (100 ng/mL, Peprotech #100-26). To generate air-liquid-interface (ALI) 2D cultures, cells were dissociated with TrypLE express enzyme and seeded onto PureCol-coated (30 µg/mL, Advanced BioMatrix)-6.5 mm transwell inserts (0.4 µm pore size polyester membrane, Corning #3470) and cultured in submerged conditions in BC expansion medium. When confluent, cells were differentiated in ALI-differentiation medium consisting of A83-01 (50 nM, Tocris #2939), hEGF (0.5 ng/mL, Peprotech #AF10015), 3,3,5-Triiodo-L-Thyronine (100 nM, Sigma #T6397), Epinephrine hydrochloride (0.5 mg/mL, Sigma #E4642), Retinioc acid receptor agonist (100 nM, Cayman #16144), Hydrocortisone (0.5 mg/mL, Sigma #H088) and Pen/Strep (1%, Gibco #15070-063) in 492,5 mL AdDMEM/F12 supplemented with additional A83-01 (500 nM, Tocris #2939). After 3-4 days of complete submerged culture conditions, the volume of medium at the apical side was decreased to a minimal amount for cells to be submerged. After 4 days, ALI-differentiation medium was removed and refreshed with ALI-differentiation medium supplemented with DAPT (5 mM, Fisher Scientific #15467109) for 14-18 days. The addition of DAPT to the differentiation medium together with the small amount of medium supplemented to the apical side of the culture shifted the differentiation program towards ciliated cells. Medium was refreshed twice a week (Mondays and Fridays), at which point the apical side was washed with PBS to remove mucus.

### Plasmids

The plasmid pCMV-Sport6-CD63-pHluorin (Addgene #130901), encoding a modified version of CD6, where the pH-sensitive fluorophore pHluorin was inserted into the first external loop, has been described before (Verweij, Bebelman et al. 2018).

### Viruses

SARS-CoV-2 strain/NL/2020 (EVAg, 010V-03903) was fully sequenced (original SARS-CoV-2 variant 19B (https://clades.nextstrain.org/) and cultivated on monkey kidney Vero E6 cells. All work with SARS-CoV-2 was conducted in Biosafety Level-3 conditions at the University Medical Center Utrecht, according to WHO guidelines (Laboratory biosafety guidance related to coronavirus disease (COVID-19)). The respective virus titer was determined by titration of infectious particles on Vero E6 cells (TCID50) and quantitative real-time reverse transcription-PCR (qRT-PCR) specific for the SARS-CoV-2 E gene (Corman, Landt et al. 2020).

### Infections

Vero E6 cells were seeded the day prior to infection in 96 well plates (for qPCR) or in 12-well plates on 18 mm precision coverslips (thickness No. 1.5H, #0117580, Marienfeld) and maintained in DMEM supplied with 2% FCS, 100 Units/ml penicillin, 100 µg/ml streptomycin and 2 mM L-glutamine. The viruses were inoculated at the indicated multiplicity of infection (MOI) for 2h at 37°C. After inoculation the virus was removed, the cells were rinsed with PBS and fresh medium was applied.

Differentiated HNECs were incubated with phosphate-buffered saline (PBS) for 20 min at 37°C to remove the mucus from the apical surface before inoculation with the respective viruses. The viruses were diluted in advanced DMEM/F-12 and inoculated apically at the indicated MOI for 2h at 37°C. After inoculation, the virus was removed and the apical side was carefully rinsed with PBS three times, followed by PBS removal and apical exposure to air.

### Quantitative real-time reverse transcription-PCR

qRT-PCR-based quantification of SARS-CoV-2 infection was performed on supernatants or cellular samples. For cellular samples, RNA was extracted from the samples using Trizol/Chloroform extraction followed by RNA isolation using an RNA isolation kit (Norgen). Quantitative real-time reverse transcription-PCR (qRT-PCR) specific for the SARS-CoV-2 E gene was performed with primers and probes in table 1 and analyzed using a StepOnePlusTM Real-Time PCR system.

**Table 1:**
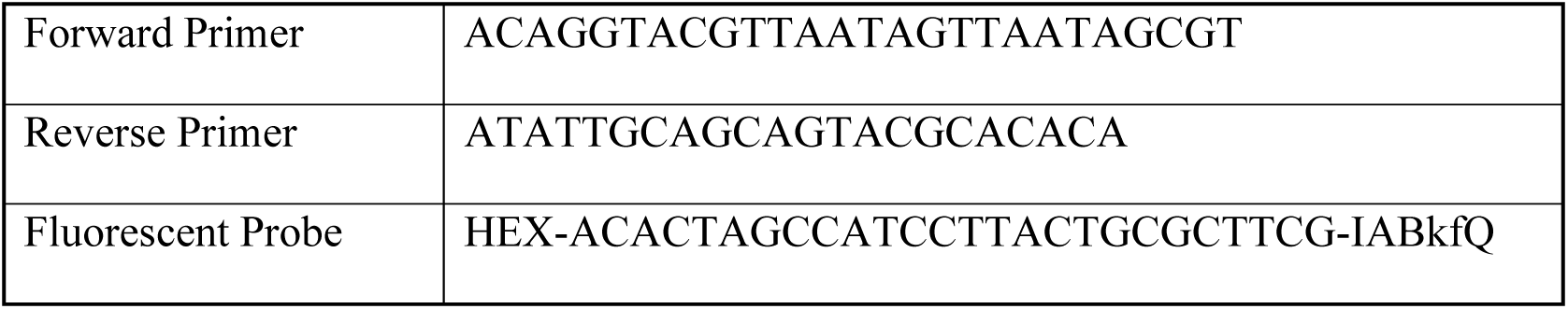
Primer and Probe sequences SARS-CoV-2 E gene.

### Antibodies

Rabbit anti-ACE2 (Sino Biological, #80031-RP02), mouse beta-tubulin IV (BioGenex, #MU178-UC), mouse anti-CC10 (Origene, AM26360PU-N), mouse anti-CD63 (DSHB Hybridoma Product H5C6, deposited to the DSHB by August, J.T. / Hildreth, J.E.K.), mouse anti-Clathrin Heavy Chain (Thermo Fisher Scientific, #MA1-065), mouse anti-dsRNA (J2, Sci Cons, #10010200), mouse anti-ERGIC53 (Enzo, #ENZ-ABS300-0100), mouse anti-GM130 (BD Biosciences, #610823), rabbit anti-Lamtor4/c7orf59 (Cell Signaling, #12284S), rabbit anti-LC3 (MBL, #PM036), mouse anti-M6PR (Abcam, ab198294), mouse anti-MUC5AC (Thermo Fisher Scientific, #MA1-38223), rabbit anti-MUC5AC (Abcam, ab198294), rabbit anti-p63 (Abcam, ab124762), rabbit anti-Rab7 (Cell Signaling, #D95F2), mouse anti-Rab27 (BD Biosciences, # 558532), mouse anti-SARS-CoV-2 spike S1 (Invitrogen, # MA5-36245), rabbit anti-SARS-CoV-2 spike S1 (Sino Biological, #40150-R007), rabbit anti-TGN46 (Merck Millipore, #ABT95), and rabbit anti-TMPRSS2 (Atlas Antibodies, Cat# HPA035787). Secondary antibodies were highly cross-adsorbed Alexa Fluor -488, -594, or -647 -conjugated goat antibodies against rabbit IgG or mouse IgG, IgG1 or IgG2A (Thermo Fisher Scientific), goat-anti-mouse and goat anti-rabbit Abberior STAR 635P and goat anti-mouse Atto 647N (Sigma-Aldrich), together with Alexa Fluor Plus 405 Phalloidin (Thermo Fisher Scientific), or Atto 647N Phalloidin (Sigma-Aldrich) and DAPI (Sigma-Aldrich).

### Immunofluorescent staining of fixed samples

Prior to fixation, HNECs were apically washed with PBS. Vero E6 cells and HNECs were fixed in 4% paraformaldehyde and 4% sucrose in PBS (w/v, w/v) for 15 min at 37°C or 20 min with prewarmed fixative for infected samples, permeabilized in 0.5% Triton X-100 in PBS (v/v) for 15 min and blocked in 3% BSA in PBS (w/v) for 30 min. Before staining of HNECs, filters were cut out from transwell chambers and sectioned into quarters. HNECs on filter fragments were suspended in a 30 µl droplet of primary antibodies diluted in 3% BSA in PBS (w/v) overnight at 4°C. Then, samples were washed three times in 0.1% Triton X-100 in PBS (v/v) for 20 min before the addition of corresponding secondary antibodies Alexa Fluor 488-, Alexa Fluor 594, Atto 647N, or Abberior STAR635p-conjugated anti-rabbit and anti-mouse together with DAPI and Phalloidin-Atto-647N (1:200) for 2 h at room temperature. Cells were washed three times with 0.1% Triton X-100 in PBS (v/v) for 10 min. When staining dsRNA, all buffers were supplemented with RNAse inhibitor (RNasin, #N2111, Promega, 1:320.000). Cells were mounted in Prolong Diamond (Thermo Scientific, #P36970), and for airway cultures, covered with precision 13 mm glass coverslips (thickness No. 1.5H, #0117580, Marienfeld).

### TREx protocol, mCLING treatment and general protein stains

For Ten-fold Robust Expansion Microscopy (TREx), cells were either fixed for 20 min with pre-warmed (37°C) 4% paraformaldehyde for specific antibody labeling in combination with general protein stains or 4% paraformaldehyde (w/v) and 0.1% glutaraldehyde (w/v) in PBS for visualization of lipid membranes. For mCLING treatment, cells were washed twice in PBS after fixation and incubated in 5 μM mCLING-Atto647N (Synaptic Systems, 710 006AT1) in PBS overnight at room temperature. The following day, cells were fixed a second time with pre-warmed (37°C) 4% paraformaldehyde (w/v) and 0.1% glutaraldehyde (w/v) in PBS. Next, cells were washed with PBS and permeabilized using 0.2% Triton X-100 in PBS (v/v) for at least 15 min. Epitope blocking and antibody labeling steps were performed in 3% BSA in PBS (w/v).

TREx was performed as described earlier (Damstra, Mohar et al. 2022, van Grinsven, Katrukha et al. 2025). In brief, samples were post-fixed with 0.1 mg/mL acryloyl X-SE (AcX; Thermo Fisher, A20770) in PBS overnight at room temperature. For gelation, monomer solution was prepared containing 1.085 M sodium acrylate (Sigma-Aldrich, 408220), 2.015 M acrylamide (AA; Sigma-Aldrich, A4058) and 0.009% N,N’-methylenebisacrylamide (BIS; Sigma-Aldrich, M1533) in PBS. Gelation of the monomer solution was initiated with 0.15% ammonium persulfate (APS) and 1.5% tetramethylethylenediamine (TEMED). To expand filter-cultured HNECs, 80 μl of gelation solution was pipetted directly in the Transwell chamber set down on a parafilm-covered glass slide. The sample was directly transferred to a 37°C incubator for 1 h to fully polymerize the gel without closing the gelation chamber. All gels, except for samples that were processed for subsequent general protein staining, were transferred to a 12-well plate and digested in TAE buffer (containing 40 mM Tris, 20 mM acetic acid and 1 mM EDTA) supplemented with 0.5% Triton X-100, 0.8 M guanidine-HCl, 7.5 U/mL Proteinase-K (Thermo Fisher, EO0491) and DAPI for 4 h at 37°C. The gel was transferred to a Petri dish, water was exchanged twice after 30 min and the sample was left expand overnight in ultrapure water.

For the general protein stains, gels were washed after polymerization in PBS for 15 min at least twice. Next, the gels were incubated for 1 h on a shaker at room temperature with either with 20 μg/mL Atto 594 NHS ester (Sigma-Aldrich, 08471) or 20 μg/mL Atto 647N maleimide (Sigma-Aldrich, 41784) in PBS, both prepared from 20 mg/mL stock solutions in DMSO, to label lysines and cysteines, respectively. These stainings highlighted various subcellular structures to different degrees, which motivated our choice for one or the other in specific experiments. For example, a NHS ester strongly labeled basal bodies, whereas maleimide nicely labeled membranous organelles. After staining, gels were washed with an excess of PBS, subsequently digested for 4 h at 37°C as described above, transferred to a Petri dish and expanded overnight. Prior to imaging, the gels were trimmed and mounted.

### Expanded sample imaging

Expanded gels were imaged using the same Leica TCS SP8 STED 3X microscope pulsed (80 MHz) with 405 nm and pulsed (80 MHz) white-light lasers, PMT and HyD detectors and spectroscopic detection with a HC PL APO 86x/1.20W motCORR STED (Leica 15506333) water objective.

### TREx data analysis

Processing and analysis of data from expanded samples was done using ImageJ and Arivis Vision4D (Arivis AG) for 3D visualization and rendering. All single planes shown in this manuscript are sum projections of 3 planes (z-spacing 0.36 or 0.15 μm), expect when volumetric rendering is indicated. Contrast was adjusted manually. Figure S2E shows a volumetrically rendered stack depth-coded in grayscale along the z-axis using Arivis to provide contrast reminiscent of scanning electron microscopy. All volumetric renders are running sum projections of 3 planes generated using the RunningZProjector plugin (https://valelab4.ucsf.edu/~nstuurman/IJplugins/Running_ZProjector.html) for ImageJ after which the data was imported into Arivis. For Figure S2D gamma was adjusted manually to increase visibility of both cilia and intracellular structures. Some rendered data was filtered in Arivis using the Discrete Gaussian Filter with smoothing radius of 2 to aid visibility.

For Figure 1D, 1E, 1G and 1H, raw data was imported into ImageJ. Acquisitions were processed using a 3D gaussian blur with a sigma value x, y and z of 0.5 or 2 for the maleimide channel and Spike/GM130/DAPI channels, respectively. Indicated planes are sum projections of 3 planes (z-spacing 0.36 μm) of the filtered stack. For all images, the maleimide channel is shown in inverted contrast. To visualize GM130 and Spike in combination with the inverted maleimide channel, multicolor images of maleimide with GM130 and Spike were presented as overlays instead of merges. In brief, manual intensity-based thresholding of GM130 and Spike channels was used to generate binary masks of these respective channels, which were then converted to regions of interest (ROI). The merged images depict the non-thresholded Spike and GM130 images, merged with the maleimide, but the maleimide signal is not shown within these GM130 and Spike-positive ROIs. For Figure 1I, virus-induced large organelles were defined as discrete bulbous organelles with internal structure that were morphologically distinct from other cellular structures. The diameter corresponds to the largest width of the structure.

For segmentation in Figure S2F (right panel), the raw dataset was imported in ImageJ and was segmented for microvilli and cilia using the trainable Weka segmentation plugin in ImageJ (Arganda-Carreras, Kaynig et al. 2017).

### Confocal and STED imaging of non-expanded samples

Confocal micrographs from non-expanded samples were acquired using a Zeiss LSM880 Fast Airyscan microscope with 405 nm, 561 nm, 633 nm and argon multiline lasers, internal Zeiss spectral 3 PMT detector and spectroscopic detection using a Plan-Apochromat 63x/1.2 glycerol or an Alpha Plan-APO 100x/1.46 Oil objective (Figures 2K, 3G).

Data from non-expanded samples shown in Figures S1A, S1B and S2F were acquired using a Leica SP8 STED 3X microscope with 405 nm and pulsed (80 MHz) white-light lasers, PMT and HyD detectors and spectroscopic detection using a HC PL APO 93x/1.3 GLYC motCORR STED (Leica 11506417) glycerol objective. For STED imaging of Alexa Fluor 594, Atto674N and Abberior STAR635p, we used 594 nm and 633 nm laser lines for excitation and a 775 nm synchronized pulsed laser for depletion. For Alexa 488 we used 488 nm excitation and a 592 nm continuous depletion laser line. For multicolor STED imaging, each fluorescent channel was imaged using the 2D STED configuration in sequential z-stack mode from highest to lower wavelength to prevent photobleaching. The z-stacks were subjected to mild deconvolution using Huygens Professional software version 17.04 (Scientific Volume Imaging, The Netherlands) with the classic maximum likelihood estimation (CMLE) algorithm and the signal-to-noise ratio (SNR) parameter equal to 7 over 10 iterations.

### Processing and analysis of confocal micrographs

Confocal micrographs were deconvolved (except for images shown in Figure 1B) in SVI Huygens (Huygens professional v21.04) and further processed and analyzed in ImageJ. If necessary, image drift correction was performed using the MultistackReg plugin for ImageJ. Cell segmentation was done manually based on phalloidin staining to generate whole cell ROIs. All quantifications in Figures 2 and S4 were obtained from summed intensity projections of the entire cell. Cellular intensities of viral infection markers dsRNA and SARS-CoV-2 Spike were calculated from deconvolved, summed, but unprocessed images. To compare individual cells across replicates, cellular infection marker intensities (defined as [(mean cellular intensity - mean extracellular background intensity) x cellular area]) were normalized for every replicate by dividing individual cellular intensities by the average cellular intensities of the mock condition. Cells from the infected condition were considered infected and included in the analysis if the cellular Spike intensity exceeded a cutoff defined as the average intensity plus four times the standard deviation of mock cells.

### Quantification of organellar marker intensities

To measure the intensity of organelle markers (GM130, TGN46, CD63, LC3, and LAMTOR4) on their respective organelles, summed intensity projections of GM130 (for Golgi), CD63, LC3, or LAMTOR were thresholded to generate ROIs for each cell. The following image processing steps were applied: Gaussian blur, morphological filtering (using a disk element of 2 pixels, MorpholibJ plugin for ImageJ (Legland, Arganda-Carreras et al. 2016)) and background subtraction (rolling ball radius of 20 pixels) for Golgi markers, and Gaussian blur, background subtraction (rolling ball radius of 10 pixels), and morphological filtering (using a disk element of 1 pixel) for CD63, LC3, and LAMTOR4 markers, followed by intensity-based thresholding. Cellular organellar ROIs were used to calculate the normalized organellar intensities of the respective organelle markers in the deconvolved, summed, but unprocessed images. To compare individual cells across different replicates, the organelle-localized intensities (defined as [(mean organellar intensity per cell-(mean cellular non-organellar intensity) x organellar area]) were normalized by dividing individual organelle-localized intensities by the average organelle-localized intensities observed in the mock condition. Additionally, Golgi fragmentation was assessed using cellular Golgi ROIs, with fragmentation defined as the ratio of convex hull area to particle area.

### Colocalization analysis

For pixel intensity cross correlation analyses, images were subjected to background subtraction (rolling ball radius of 50 pixels) before calculating the non-thresholded Pearson’s R Value using the Coloc 2 plugin for ImageJ.

### High content microscopy

Whole 13 mm coverslips were imaged with a Cell Discoverer 7 automated microscope (Zeiss) equipped using a Plan-Apochromat 5x/0.35 dry objective, a Castor 2x tubelens in the lightpath (effective magnification 10x), 385 nm, 567 nm and 590 nm LEDs for excitation and a Axiocam 712 mono camera, controlled by Zen blue software (Zeiss).

### Image-based toxicity assay

Vero E6 cells were seeded on 13 mm coverslips at a density of 12.5%. 16 h after seeding, cells were either fixed or treated with Nexinhib20 in a concentration range of 128 to 2000 nM, or treated with DMSO as a control. Subsequently, cells were fixed at 2, 24, or 48 h after treatment. Nuclei were stained with DAPI, and coverslips were mounted in Prolong Diamond. Imaging of cells was performed using a Zeiss Cell Discoverer 7 automated microscope, as described above. Nuclei were detected and counted using the Stardist plugin for ImageJ (Schmidt 2018), within the entire coverslip excluding any damaged or aberrant regions.

### Image-based viral proliferation assay

To determine the spread of SARS-CoV-2 infection by immunofluorescent imaging, 100.000 Vero E6 cells were seeded on 13 mm coverslips. The following day, cells were infected with SARS-CoV-2 at MOI 0.01. 2 h post infection, coverslips were transferred to new medium supplemented with Nexinhib20 (128-2000 nM) or DMSO and fixed after 24 h or 48 h. Cells were stained for SARS-CoV-2 Spike and CD63 and coverslips were mounted in Prolong Diamond.

Imaging of cells was performed using a Zeiss Cell Discoverer 7 automated microscope, as described above. A custom whole cell segmentation model was trained in Cellpose 2.0 (Pachitariu and Stringer 2022), using user-guided segmentation of the CD63 staining. This trained model was then employed to segment full entire coverslips. Next, raw images were processed in FIJI. A background subtraction was performed (50-pixel rolling ball), after which cellular intensities of SARS-CoV-2 Spike and CD63 were measured within a ROI that excluded areas outside the coverslip as well as debris on the coverslip, using Cellpose segmentation. Next, the Spike/CD63 ratio was calculated (defined as [(mean Spike intensity - mean Spike extracellular background intensity) x cellular area) / (mean CD63 intensity - mean CD63 extracellular background intensity) x cellular area)]). Cells were classified as infected if their Spike/CD63 ratio exceeded the mean plus four times the standard deviation of the uninfected control. Subsequently, the fraction of infected cells was determined by dividing the number of infected cells by the total number of cells analyzed. To provide a comparative analysis, the fraction of infected cells was normalized to the condition with the highest number of infected cells, which corresponded to untreated infected cells at 2 dpi. Finally, the normalized data was averaged across three independent experiments to obtain the final results.

### TIRF imaging of CD63 secretion

Secretion of CD63^+^ endolysosomes was analyzed by TIRF imaging of CD63-pHluorin-expressing Vero E6 cells (Verweij, Bebelman et al. 2018). In brief, cells were seeded on 24 mm coverslips and transfected with a plasmid encoding a version of CD63 in which the pH sensitive fluorophore pHluorin had been inserted into the first external loop. 24 h after transfection, cells on coverslips were mounted in an Attofluor imaging ring (Thermo Fisher Scientific # A7816) and in 37°C prewarmed medium, supplemented with nexinhib20 (1-2 µM) or DMSO. After 1 h pre-incubation, CD63-pHluorin was imaged at 3 frames/s using an objective-based azimuthal ILAS2 TIRF microscope (Nikon Eclipse Ti, modified by Roper Scientific), equipped with Nikon Apo TIRF 100x N.A. 1.49 oil objective, a Photometrics Prime BSI camera, 100 mW 561 nm laser and 49008 Chroma filter set, and controlled with MetaMorph 7.8.8 software (Molecular Device). Fusion events were detected as sudden increases in fluorescent intensity and were quantified as the number of events per cell scored per minute for the duration of imaging (3 min).

## ACKNOWLEDGEMENTS

We thank Nalan Liv for generously providing antibodies and members of the Beekman lab for providing assistance in setting up HNEC cultures at Utrecht University. This work was supported by the Eindhoven-Wageningen-Utrecht Alliance (www.ewuu.nl) that supports the Centre for Living Technologies. The authors declare no competing financial interests. Additional funding came from the PPP allowance made available by Health∼Holland, Top Sector Life Sciences & Health to stimulated public-private partnerships. The authors declare no competing financial interests.

## SUPPLEMENTAL FIGURES

**Supplemental Figure 1, related to Figure 1.**
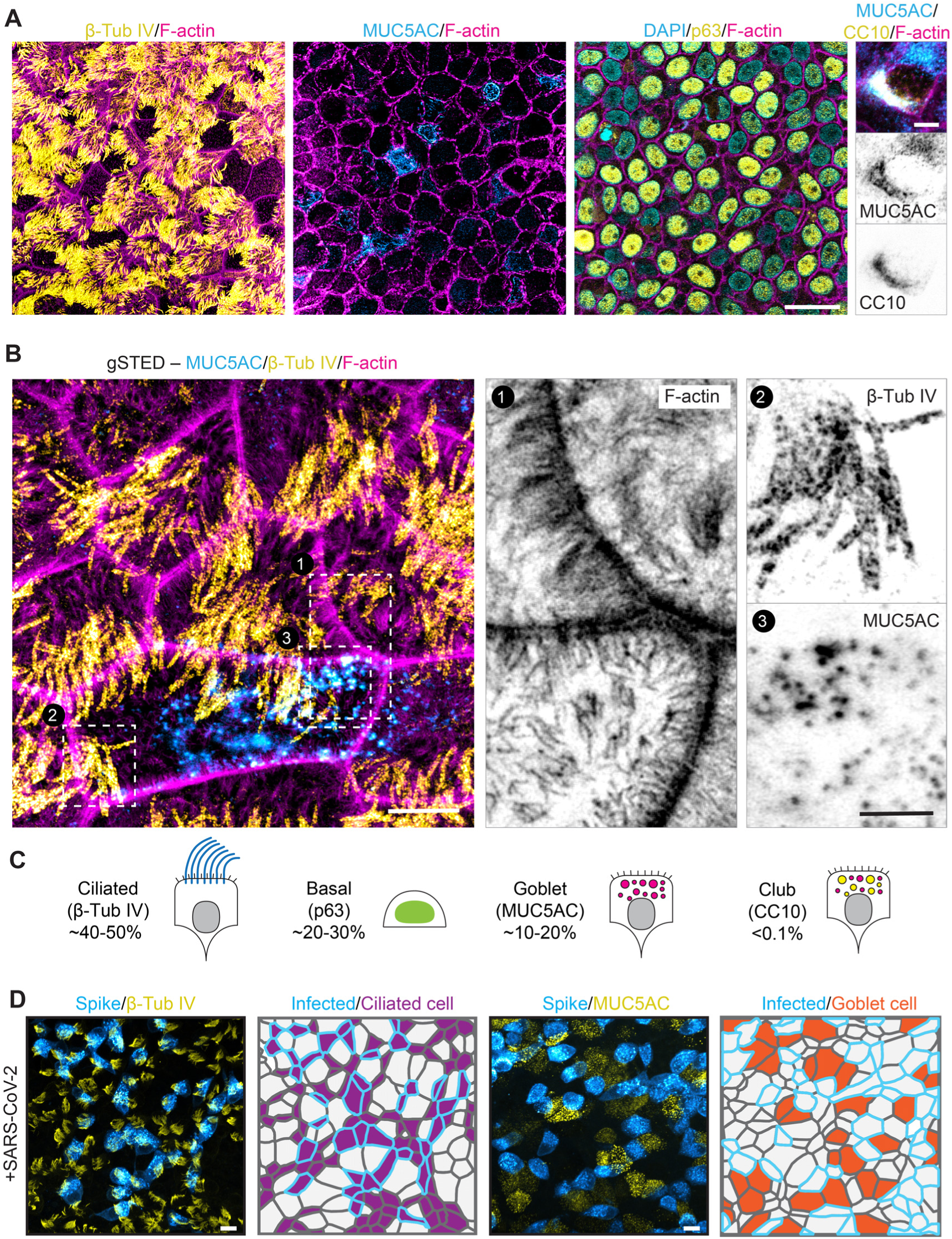
Characterization of human airway cultures. (A) Validation of differentiated airway cultures used in this study by immunofluorescent imaging of markers for distinct cell types (from left to right): β-IV Tubulin (β-Tub IV) (multiciliated cells, yellow), MUC5AC (goblet cells, blue), p63 (basal cells, yellow) and a combination of MUC5AC (blue) and CC10 (club cells, yellow). All images are co-stained with F-actin (phalloidin, magenta). (B) gated STED (gSTED) image of human nasal epithelial cells (HNECs) stained for F-actin (magenta) (1), β-Tub IV (yellow) (2) and MUC5AC (blue) (3). (C) Quantification of percentage of indicated cell types from samples as in (A). (D) Immunofluorescent image and segmentation of infected HNECs stained for spike (blue) and β-Tub IV (multiciliated cells, yellow, left) or MUC5AC (goblet cells, yellow, right), showing preferential infection of multiciliated cells. All SARS-CoV-2 infections were performed at Multiplicity of Infection (MOI): 0.1, and cells were fixed at 3 days post infection (dpi). Scale bars are 20 µm (A overview), 10 µm (D), 5 µm (A zoom and B overview), or 2 µm (Zoom B).

**Supplemental Figure 2, related to Figure 1:**
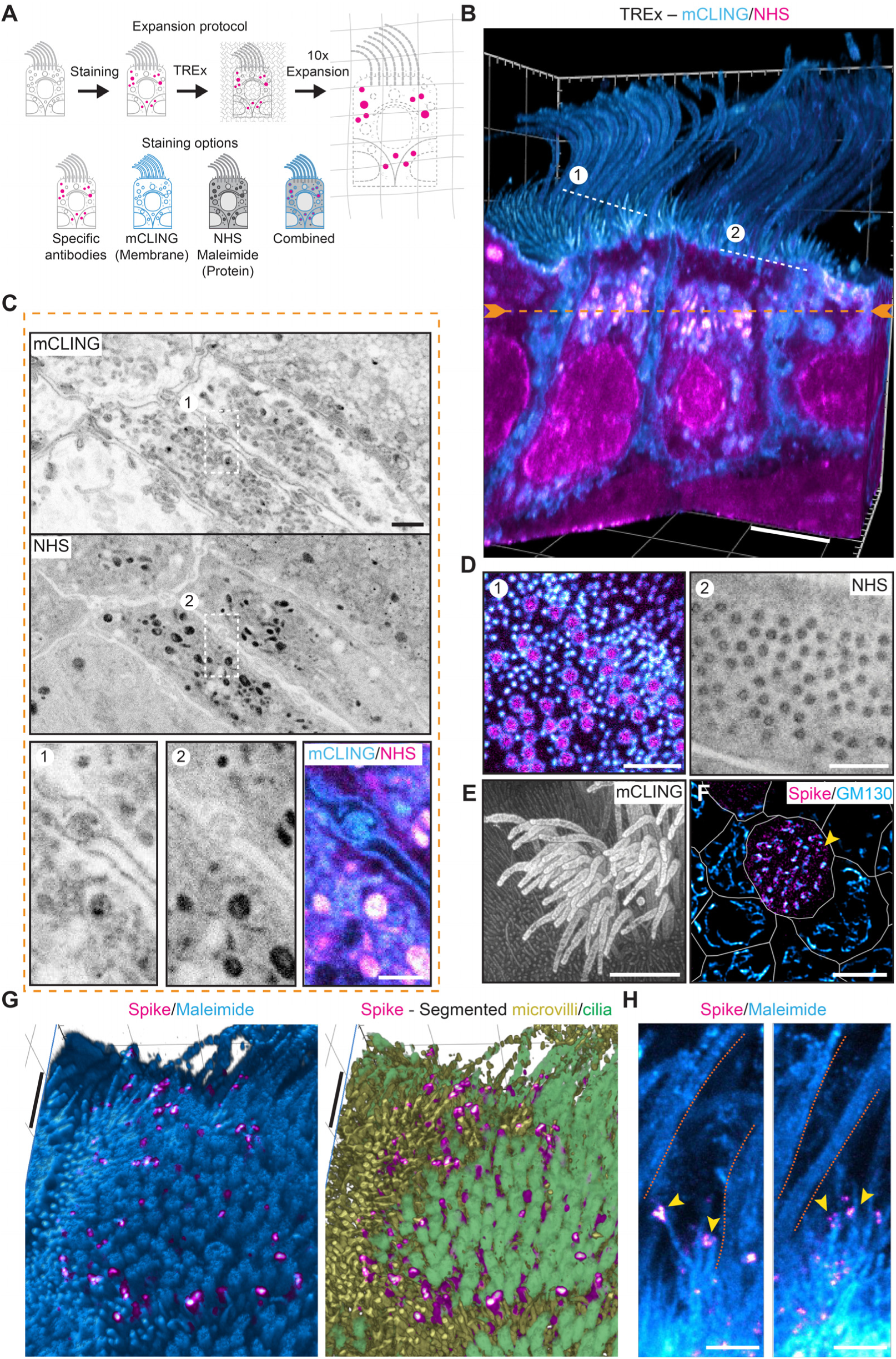
TREx microscopy reveals ultrastructure of human airway cells. (A) Ten-fold Robust Expansion (TREx) microscopy principle. Human nasal epithelial cells (HNECs) are stained with specific antibodies or small molecule stains (mCling, NHS, maleimide) and embedded in a swellable polymer gel that isotropically expands ten-fold after proteolytic digestion. (B-E) Volumetric 3D renders of TREx of HNECs stained for mCLING (total membrane) and a NHS-ester (total protein), highlighting different aspects of cellular morphology for the same cellular volume. See also movie S1. (B) Volumetric 3Drender of multiciliated cell stained for mCLING (blue) and a NHS-ester (magenta) clipped to reveal intracellular ultrastructure. Dotted orange and white lines mark planes shown in C and D, respectively. (C) Single plane from B. Zoom shows elaborate interdigitated membrane contacts between cells. (D) Single planes from (B) showing basal bodies with mCLING (blue) and a NHS-ester (magenta) (D-1) or a NHS-ester (D-2). (E) Top-down depth-encoded volumetric 3D render of multiciliated cell shown in (B) providing contrast reminiscent of scanning electron microscopy. (F) gated STED (gSTED) image of SARS-CoV-2 infected multiciliated cells stained for spike (magenta) and GM130 (Golgi, blue). White lines show segmented cells, yellow arrow indicates an infected cell with fragmented Golgi. (G) Volumetric 3D render of TREx image of an SARS-CoV-2 infected multiciliated cell stained for maleimide (blue) and spike (magenta) showing Spike puncta colocalizing on apical membrane protrusions. Right image shows corresponding segmentation of microvilli (yellow) and cilia (green) (left). (H) Single planes TREx image of SARS-CoV-2 infected multiciliated cell stained for maleimide (blue) and Spike (magenta) showing spike-positive apical protrusions. Orange lines mark cilia, yellow arrows point to Spike puncta. Scale bars are 500 nm (G). Scale bars (corrected to indicate pre-expansion dimensions): B, E ∼2 μm, C, D ∼1 μm, G and zooms ∼500 nm. TREx scale bar was corrected to indicate pre-expansion dimensions. All SARS-CoV-2 infections were performed at Multiplicity of Infection (MOI): 0.1, and cells were fixed at 3 days post infection (dpi).

**Supplemental Figure 3, related to Figure 2.**
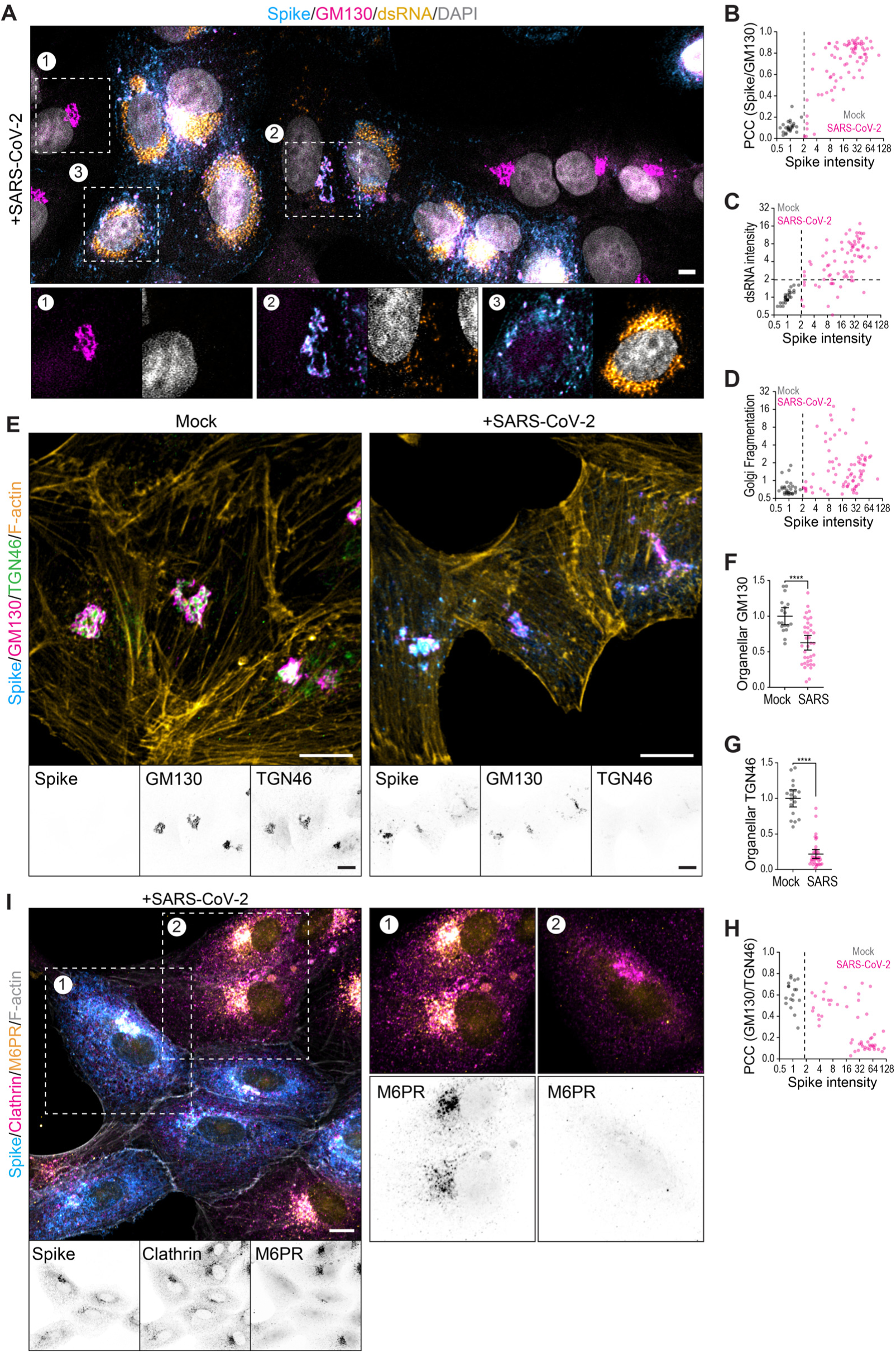
Quantitative analysis of the SARS-CoV-2 infection in Vero cells. (A) Confocal images of uninfected and SARS-CoV-2 infected Vero E6 cells stained for spike (blue), GM130 (cis-Golgi, magenta), double stranded (ds) RNA (yellow) and DAPI (grey). Boxes mark cells with progressive increase of viral markers and disruption of Golgi morphology (from 1 to 3). (B-D) Comparison between uninfected (Mock) and SARS-CoV-2 infected cells. Quantification of the colocalization of spike and GM130 (non-thresholded Pearsons correlation coefficient (PCC)) relative to the normalized cellular Spike intensity (on a 2 log scale) (B); the normalized cellular dsRNA intensity (C) and the fragmentation of the Golgi (D) relative to the cellular Spike intensity. n=number of analyzed cells: n= 26-67 from 3 independent replicates (refer to supplemental table 1 for details). (E) Confocal images of uninfected (Mock) and SARS-CoV-2 infected (+SARS-CoV-2) Vero E6 cells stained for spike (blue), GM130 (magenta), TGN46 (trans-Golgi, green) and F-actin (phalloidin, yellow). (F-H) Comparison between uninfected (Mock) and SARS-CoV-2 infected cells. Quantification of the normalized organelle marker intensity (calculated as (mean organellar intensity – mean non organellar intensity) x organelle area) per cell of GM130 (F) and TGN46 (G). Graph H shows the colocalization of GM130 and TGN46 in relation to the cellular Spike intensity, similar to graph B. n=number of analyzed cells: n= 18-37 from 3 independent replicates (refer to supplemental table 1 for details). (I) Immunofluorescent confocal imaging of SARS-CoV-2 infected Vero E6 cells stained for spike (blue), Clathrin Heavy Chain (Clathrin, magenta), mannose-6-phosphate receptor (M6PR, magenta) and F-actin (phalloidin, grey) The quantified data are derived three independent experiments (see supplemental table 1). Dots represent individual cells, bars indicate mean±SD. **** p<0.0001 (Student’s *t*-test, unpaired). Scale bars are 10 µm. All SARS-CoV-2 infections were performed at Multiplicity of Infection (MOI): 0.01, and cells were fixed at 2 days post infection (dpi).

**Supplemental Figure 4, related to Figure 2.**
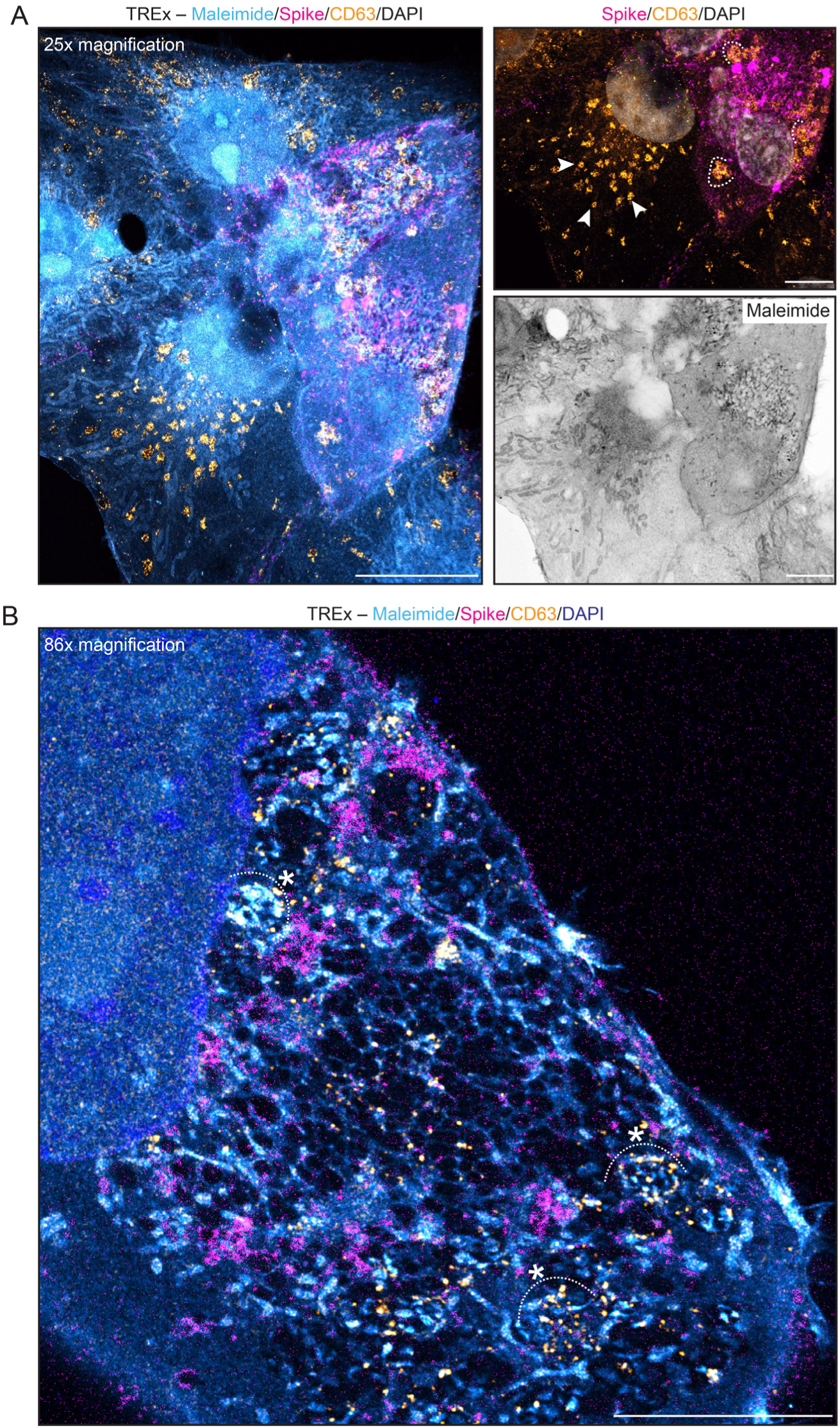
Multi-scale TREx imaging of SARS-CoV-2 infected Vero E6 cells. (A) Maximum projection of TREx image of SARS-CoV-2 infected Vero E6 cell stained for maleimide (blue), spike (magenta), CD63 (yellow) and nuclei (DAPI, dark blue). Arrows indicate small CD63-positive vesicles in Spike-negative cell and curved lines indicate enlarged Spike-positive, CD63-positive vesicles in infected cell. (B) TREx image similar to (A) with a more detailed example of the spike-positive, CD63-positive enlarged vesicles, indicated by curved lines with asterisk. Scale bars are 10 µm (A), 5 μm (Zooms in A). TREx scale bars were corrected to indicate pre-expansion dimensions. All SARS-CoV-2 infections were performed at Multiplicity of Infection (MOI): 0.01, and cells were fixed at 2 days post infection (dpi).

## SUPPLEMENTAL VIDEO LEGENDS

**Supplemental movie 1:**

This video corresponds to Supplemental Figure 2B-E. TREx imaging of human airway cells stained with membrane (blue) and protein (magenta) dyes. Various clipping planes are used and single sections are presented to show the utility of this technique in visualizing complex structures, especially through the use of total membrane and protein stains to provide structural context.

**Supplemental movie 2:**

This video corresponds to Figure 1F. TREx imaging of human airway cells infected with SARS-CoV-2 stained for total proteins (inverted greyscale) and spike (magenta). Orange arrowheads indicate virus-induced intracellular structures. Circles and arrowheads indicate virus-induced large organelles

**Supplemental movie 3:**

This video corresponds to Figure 2L. TREx imaging of Vero E6 cells infected with SARS-CoV-2 stained for total proteins (blue), spike (magenta) and CD63 (yellow/orange). Large Spike densities are apparent, as well as enlarged vesicles. The external surface of these vesicles is highlighted in green, and these structures are subsequently shown in isolation. A virus-containing enlarged vesicle that appears to fuse with the plasma membrane is circled. Here, arrowheads mark intralumenal vesicles visible from the outside of the cell.

**Supplemental movie 4:**

This video corresponds to Figure 3B. TIRF imaging of CD63-pHluorin secretion in Vero E6 cells treated with Nexinhib20 or DMSO control.

